# Male rats emit aversive 44-kHz ultrasonic vocalizations during prolonged Pavlovian fear conditioning

**DOI:** 10.1101/2023.04.06.535936

**Authors:** Krzysztof H. Olszyński, Rafał Polowy, Agnieszka D. Wardak, Izabela A. Łaska, Aneta W. Grymanowska, Wojciech Puławski, Olga Gawryś, Michał Koliński, Robert K. Filipkowski

## Abstract

Rats are believed to communicate their emotional state by emitting two distinct types of ultrasonic vocalizations. The first is long “22-kHz” vocalizations (>300 ms, <32 kHz) with constant frequency, signaling aversive states and the second, is short “50-kHz” calls (<150 ms, >32 kHz), often frequency-modulated, in appetitive situations. Here we describe aversive vocalizations emitted at a higher pitch by male Wistar and spontaneously hypertensive rats (SHR) in an intensified aversive state – prolonged fear conditioning. These calls, which we named “44-kHz” vocalizations, are long (>150 ms), generally at a constant frequency (usually within 35-50 kHz range) and have an overall spectrographic image similar to 22-kHz calls. Some 44-kHz vocalizations are comprised of both 22-kHz-like and 44-kHz-like elements. Furthermore, two separate clustering methods confirmed that these 44-kHz calls can be separated from other vocalizations. We observed 44-kHz calls to be associated with freezing behavior during fear conditioning training, during which they constituted up to 19.4% of all calls and most of them appeared next to each other forming uniform groups of vocalizations (bouts). We also show that some of rats’ responses to the playback of 44-kHz calls were more akin to that of aversive calls, e.g., heart rate changes, whereas other responses were at an intermediate level between aversive and appetitive calls. Our results suggest that rats have a wider vocal repertoire than previously believed, and current definitions of major call types may require reevaluation. We hope that future investigations of 44-kHz calls in rat models of human diseases will contribute to expanding our understanding and therapeutic strategies related to human psychiatric conditions.

## Introduction

Charles Darwin wrote: “That the pitch of the voice bears some relation to certain states of feeling is tolerably clear” (Darwin, 1872). This has also been tolerably clearly observed and widely described for ultrasonic vocalizations of rats (Brudzynski, 2019, Brudzynski, 2021, Simola and Granon, 2019) which emit low-pitched aversive calls and high-pitched appetitive calls. The former are “22-kHz” vocalizations (Figs 1A, 2A), with 18 to 32 kHz frequency range, monotonous and long, usually >300 ms, and are uttered in distress (Brudzynski, 2013, Brudzynski, 2019, Brudzynski, 2021, Simola and Granon, 2019). The latter are “50-kHz” vocalizations (Fig. 1C), are relatively short (10-150 ms), frequency-modulated, usually within 35-80 kHz, and they signal appetitive and rewarding states (Simola and Granon, 2019, Brudzynski, 2013, Brudzynski, 2019, Brudzynski, 2021). Therefore, these two types of calls communicate the animal’s emotional state to their social group (Brudzynski, 2013). Low-pitch (<32 kHz), short (<300 ms; Fig. 1B) calls, assumed to also express a negative aversive state, have been described but their role is not clearly established (Brudzynski, 2013). Notably, high-pitch (>32 kHz), long and monotonous ultrasonic vocalizations have not yet been described. Here we show these unmodulated rat vocalizations with peak frequency^1^ at about 44 kHz (Figs 1B, 1E, 2B), emitted in aversive experimental situations, especially in prolonged fear conditioning.

**Fig. 1.**
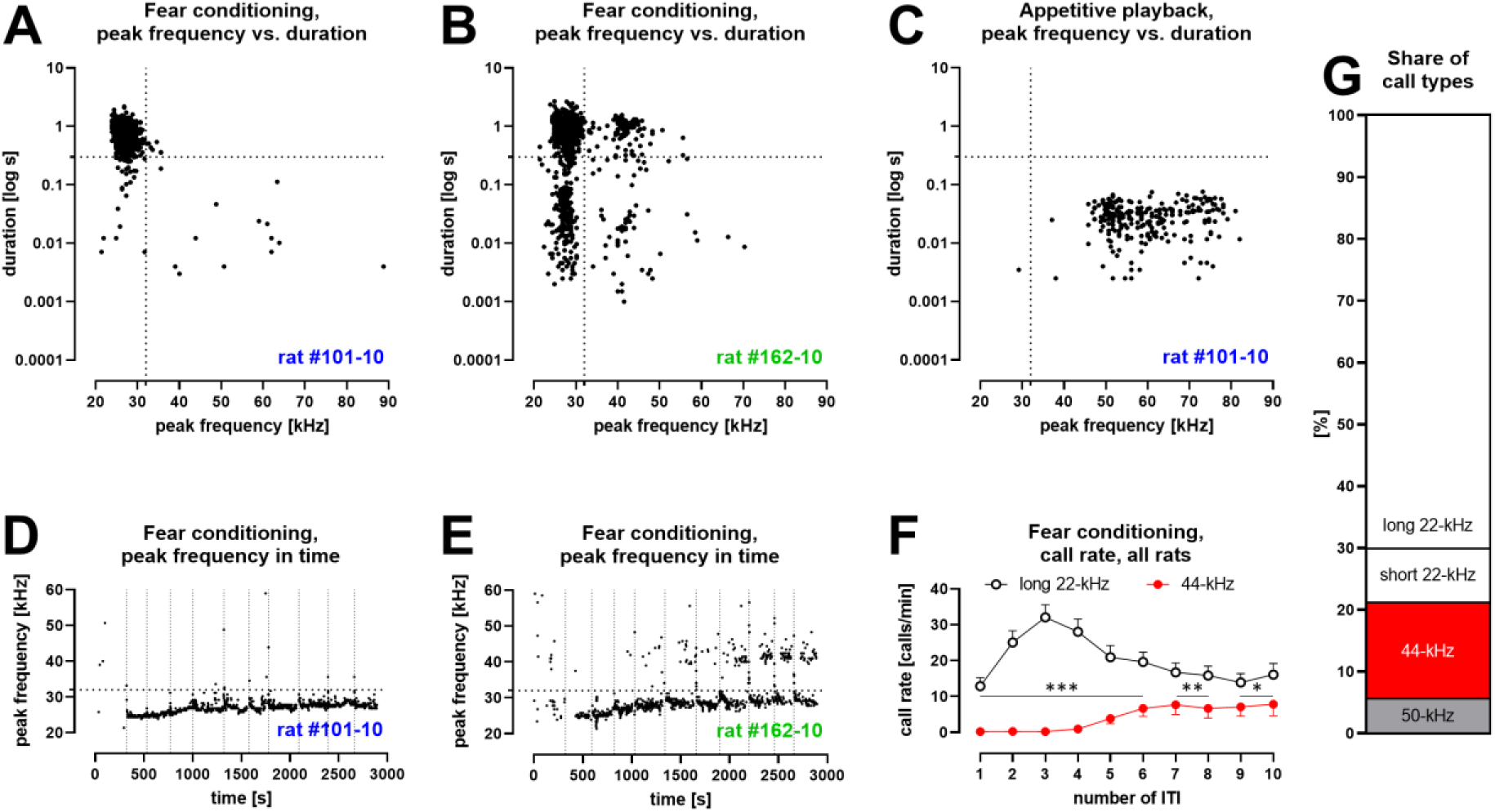
Characteristics of vocalizations emitted by Wistar rats during fear conditioning with ten aversive foot-shocks. (Tab. 1/ Exp. 1-3/#2,4,8,13; n = 46)**. A** – some rats produced aversive 22-kHz vocalizations with typical features, i.e., constant-frequency of <32 kHz, >300 ms duration – both values marked as dotted lines); example emission from one rat. **B** – some rats produced 44-kHz vocalizations with constant frequency of >32 kHz and long duration (>150 ms); example emission from one rat. **C** – rats which emitted aversive vocalizations during fear session, produced 50-kHz vocalizations during appetitive playback session the following day (full data published in Olszyński et al., 2021); representative data from same rat in **A**. **D** – the onset of long 22-kHz alarm calls typically occurred after first shock stimulus (vertical dotted lines mark time of shock deliveries in **DE**); note the gradual rise in peak frequency^a^, not exceeding 32 kHz (horizontal dotted line in **DE**); data from the same rat as **AC**. **E** – in rats that emitted 44-kHz calls, the onset was usually delayed to after several foot-shocks; note the gradual rise in peak frequency of both long 22-kHz and 44-kHz vocalizations throughout training (comp. Fig. 1S2CD); data from same rat in **B**). **F** – call rate of long 22-kHz calls was higher than 44-kHz calls (*p < 0.05, **p < 0.01, ***p < 0.001) and with different time-course – maximum number of 22-kHz calls at ITI-3 (higher than ITI-1, 2, 5-10; <0.0001–0.0005 p levels); and higher number of 44-kHz calls at ITI-5-10, i.e., 6.6 ± 2.3 vs. ITI-1-4, i.e., 0.4 ± 0.2; p < 0.0001; all Wilcoxon); numbers of ITI (inter-trial-intervals) correspond to the numbers of previous foot-shocks, values are means ± SEM. **G** – long 44-kHz vocalizations had a higher incidence rate (15.5%) than short 22-kHz (8.8%) and 50-kHz calls (5.6%); values are calculated for sum of all vocalizations obtained during entire training sessions (there were fewer 50-kHz calls, i.e., 3.7%, when vocalizations prior to the first shock were not included). **A-E**: dots reflect specified single rat values. **FG**: n = 46, other results from these rats are previously published (Olszyński et al., 2021, Olszyński et al., 2022).

**Fig. 2.**
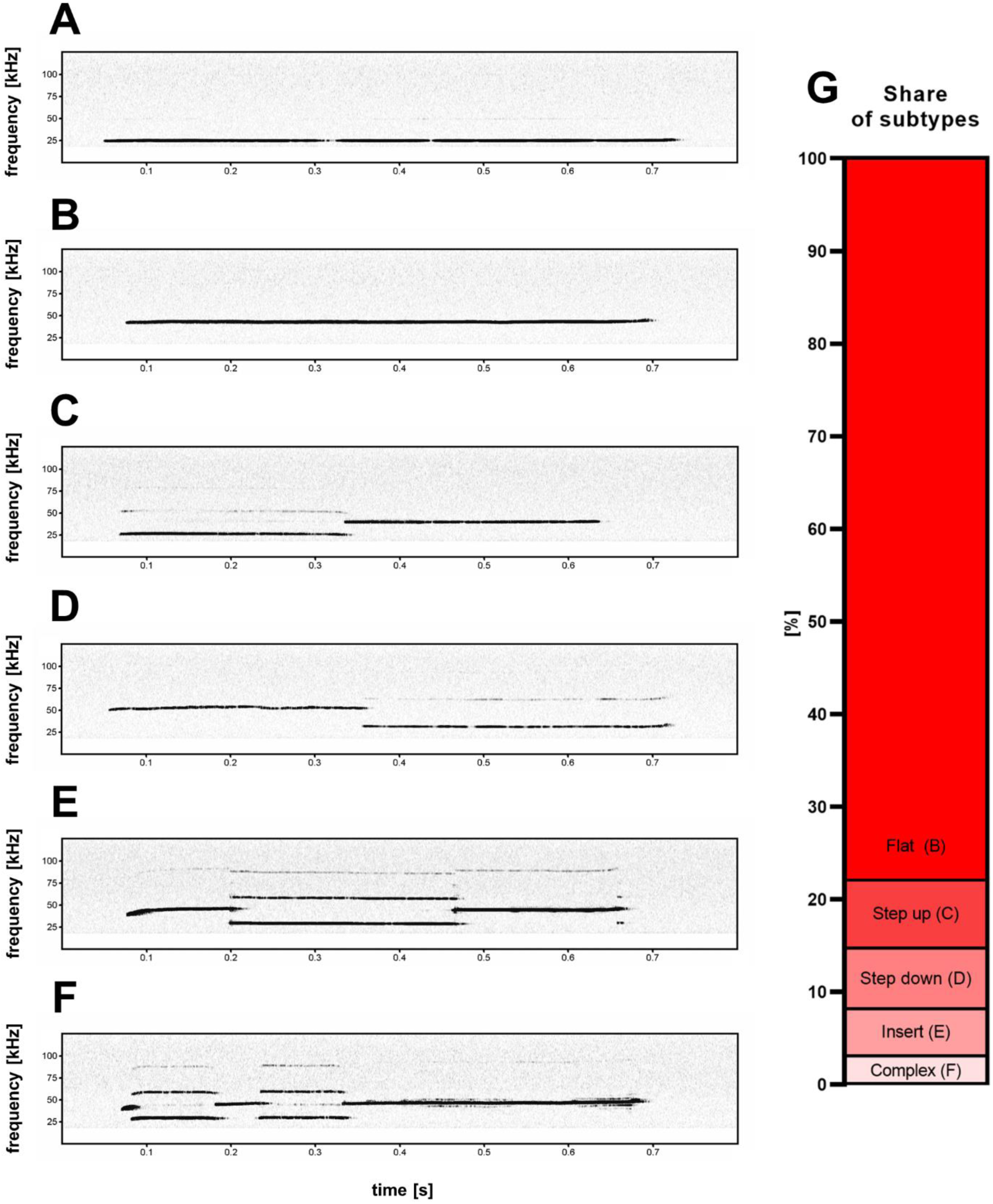
Five subtypes (B-F) of high frequency 44-kHz aversive vocalizations. **A** – standard aversive 22-kHz vocalization with peak frequency <32 kHz (peak frequency = 24.4 kHz). 44-kHz aversive vocalization subtypes: **B** – flat (constant frequency call; peak frequency = 42.4 kHz), **C** – step up (peak frequency = 39.5 kHz), **D** – step down (peak frequency = 52.2 kHz), **E** – insert (peak frequency = 38.5 kHz), **F** – complex (peak frequency = 46.3 kHz). **G** – percentage share of 44-kHz call-subtypes in all cases of detected 44-kHz vocalizations.

## Results

### High, long, and unmodulated calls

In three separate experiments (all summarized in Tab. 1/Exp.1-3, see Methods), i.e., one with trace-fear-conditioning (Tab. 1/Exp. 1) and two with delay-fear-conditioning (Tab. 1/Exp. 2-3), one of which has already been described (Tab. 1/Exp. 2, Olszyński et al., 2021, Olszyński et al., 2022), 53 of all 84 conditioned Wistar rats (Tab. 1/Exp. 1-3/#2,4,6-8,13, Figs 1B, 1E, 1S1BC) displayed vocalizations that were high-pitched, i.e., in the range of 50-kHz calls, but long and monotonous (Fig. 2B). These vocalizations, e.g., top-right group in Figs 1B and 1S1C, were outside the defined range (Brudzynski, 2019, Brudzynski, 2021, Simola and Granon, 2019) for both 50-kHz (bottom-right group in Figs 1C, 1S1A-C) and 22-kHz calls (top-left group in Figs 1A, 1B, 1S1A-C). These vocalizations were also observed in a different rat strain acquired from a different breeding colony, i.e., spontaneously hypertensive rats (SHR) (Okamoto and Aoki, 1963), also trained in delay fear conditioning (Tab. 1/Exp. 2/#10-12; Olszyński et al., 2022). Six of the 49 conditioned SHR displayed high-pitch, long, monotonous vocalizations (e.g., Fig. 2S1G); moreover, we observed more of these vocalizations in Wistar rats compared to SHR (Tab. 1/Exp. 2/#6-8,10-12) in both training, p < 0.0001, and test sessions, p = 0.0030, Mann-Whitney.

Overall, we analyzed 140,149 vocalizations from all fear conditioning experiments (Tab. 1/Exp. 1-3/#1-13, n = 218) and through trial-and-error, we set **new criteria,** namely **peak frequency of >32 kHz and >150 ms duration** to define the calls described above. We manually verified the results on the spectrogram using these parameters and only 308 calls (0.2%) were incorrectly assigned (i.e., exceptionally long 50-kHz vocalizations misplaced in the newly-defined group or borderline-short vocalizations of the newly-defined group misplaced to 50-kHz calls). Hence the new parameters correctly assigned 99.8% of cases and are thus effective to distinguish the newly-defined calls in an automated fashion. Finally, 10,445 newly-defined calls were identified, which constituted 7.5% of the total calls during fear conditioning experiments (Tab. 1/Exp. 1-3; comp. Fig. 1G). These vocalizations have a peak frequency range from 32.2 to 51.5 kHz (95% of cases) with an average peak frequency of 42.1 kHz, and they exhibited 43.8 kHz peak frequency at the cluster center in a DBSCAN analysis (Fig. 3A). In line with the accepted nomenclature convention, underlining the relationship with 22-kHz vocalizations, we christened this newly-defined calls as “**44-kHz vocalizations**”.

**Fig. 3.**
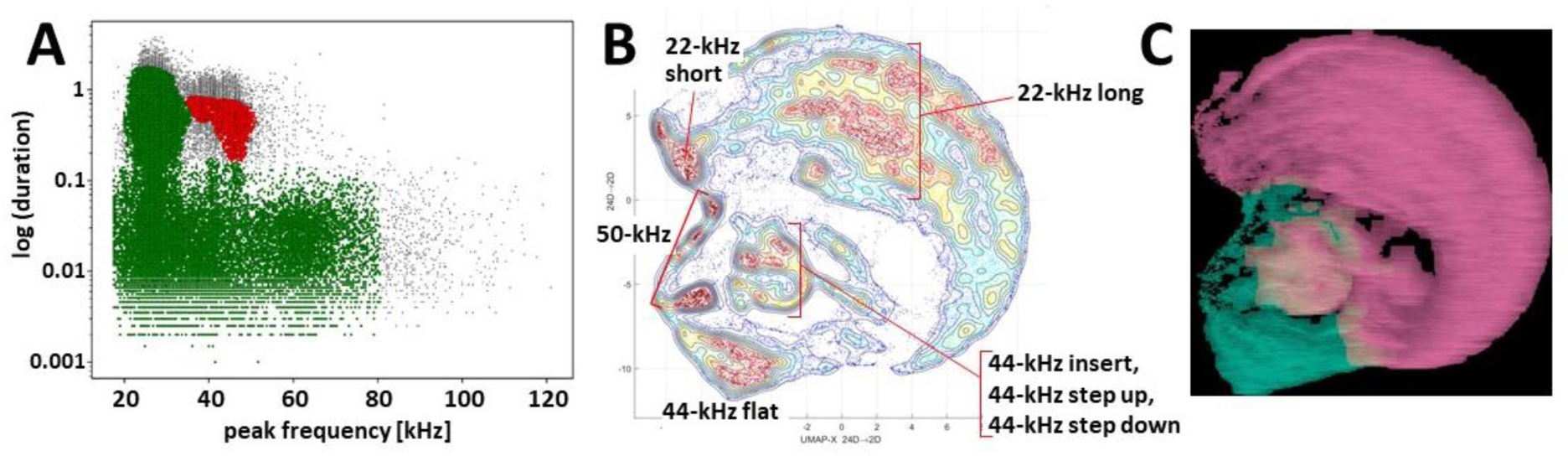
Clustering of ultrasonic vocalizations from fear conditioning sessions using two independent methods. **A** – DBSCAN algorithm (ε = 0.14) clustering of vocalizations from all fear conditioning experiments (Tab. 1/Exp. 1-3/#1-13, n = 218), silhouette coefficient = 0.198, two clusters emerge, cluster of green dots n = 77,243 (due to high generality of cluster average peak frequency and duration deemed redundant), cluster of red dots n = 5,646 (average peak frequency = 43,826.6 Hz, average duration = 0.524 s), some calls were not assigned to any cluster, i.e., outlier vocalizations, black dots, n = 4,139. **BC** – clustering by k-means algorithm and visualization of calls emitted by selected rats, i.e., with >30 of 44-kHz vocalizations, during trace and delay fear conditioning training (n = 26, selected from Tab. 1/Exp. 1-3/#2,4,7,8,11-13), total number of calls n = 40,084. **B** – topological plot of ultrasonic calls using UMAP embedding, particular agglomerations of calls labeled with their type or subtype. **C** – spectrogram images from DeepSqueak software superposed over plot B, colors denote clusters from unsupervised clustering, number of clusters set using elbow optimization (max number = 4), two clusters emerge; see also Fig. 3S1.

### 44-kHz calls in long aversive stimulation

We found 44-kHz vocalizations especially in rats which received multiple electric shocks. When we analyzed all Wistar rats that had undergone 10 trials of fear conditioning (Tab. 1/Exp. 1-3/#2,4,8,13; n = 46), these vocalizations were less frequent following the first trial (1.2 ± 0.4% of all calls), and increased in subsequent trials, particularly after the 5^th^ (8.8 ± 2.8%), through the 9^th^ (19.4 ± 5.5%, the highest value), to the 10^th^ (15.5 ± 4.9%) trials, where 44-kHz calls gradually replaced 22-kHz vocalizations in some rats (Fig. 1F, 1S2AB, Video 1; comp Fig. 1D vs. 1E). Please note, majority of the 22-kHz calls were emitted after the 3^rd^ shock, i.e., during the 3^rd^ ITI (inter-trial-interval), while 44-kHz vocalizations were emitted in the second part of the training, i.e., 5^th^ to 10^th^ ITI (Fig. 1F, comp. Fig. 1S2AB). From this group of rats (n = 46), n = 41 (89.1%) emitted long 22-kHz calls, and 32 of them (69.6%) emitted 44-kHz calls, i.e., every animal that produced 44-kHz calls also emitted long 22-kHz calls (Fig. 1S2AB). The prevalence of 44-kHz calls varied greatly among individual rats, such that for n = 3 rats, 44-kHz vocalizations accounted for >95% of all calls during at least one ITI (e.g., 140 of total 142, 222 of 231, and 263 of 265 tallied 44-kHz calls), and in n = 9 rats, 44-kHz vocalizations constituted >50% of calls in more than one ITI. The prevalence of 44-kHz calls in all experimental conditions analyzed in all animal groups is shown in Fig. 1S3.

Notably, there were more 44-kHz vocalizations during fear conditioning training than testing in all fear-conditioned Wistar rats (Tab. 1/Exp. 1-3/#2,4,6-8,13; n = 84; 3.63 ± 0.99 vs. 0.23 ± 0.13 calls/min; p < 0.0001; Wilcoxon).

In a recent publication during this paper’s review process, Gonzalez-Palomares et al. (2023), in line with the findings reported here, investigated and described 44-kHz vocalizations following prolonged (10-trial protocol) odor fear conditioning. These calls were observed predominantly during the late ITI, i.e., 8^th^-10^th^ ITI (Gonzalez-Palomares et al., 2023; Fig. S4C; please note 4^th^-7^th^ ITI were not investigated) after the shock presentations (Fig. S4B therein), which complement our results.

### Changes in frequency, duration, and mean power of long aversive calls during conditioning

Analyzing Wistar rats that undergone 10 trials of fear conditioning (Tab. 1/Exp. 1-3/#2,4,8,13; n = 46), we also observed the frequencies of 22-kHz calls to gradually rise throughout fear conditioning training, i.e., during subsequent ITI – from 24.5 ± 0.1 to 27.9 ± 0.4 kHz (Figs 1DE, 1S2C; p < 0.0001, Friedman, p = 0.0039, Wilcoxon). The frequency levels of 44-kHz vocalizations also appeared to rise – from 37.8 ± 2.1 to 39.6 ± 1.3 kHz (Figs 1E, 1S2C) but we were unable to statistically demonstrate it (p = 0.0155, Friedman, p = 0.0977, Wilcoxon).

There was a shortening of long 22-kHz calls during the first four ITI from 969.6 ± 43.1 ms to 794.6 ± 39.8 ms (p < 0.0001, Friedman; p < 0.0001, Wilcoxon, Fig. 1S2D), while 44-kHz vocalizations were longest during the 4^th^ ITI (the time of their substantial appearance, comp. Fig. 1F), i.e., 775.0 ± 135.7 ms, and shortened over subsequent ITI (619.6 ± 58.1 ms for the 10^th^ ITI, Fig. 1S2D, p = 0.0227, Friedman; p = 0.0234, Wilcoxon).

Finally, the sound mean power of 44-kHz vocalizations appeared to remain stable throughout the 10-trial sessions, while during the first half of the training, i.e., 1^st^-5^th^ ITI, 22-kHz calls were not only significantly more frequent but also louder than during the second half, i.e., 6^th^-10^th^ ITI (p < 0.0001, Wilcoxon). Consequently, long 22-kHz calls appeared louder than 44-kHz calls (p = 0.0397-0.0038, Mann-Whitney). However, in the second half of the session, this difference dissipated due to the diminishing amplitude of 22-kHz vocalizations (p = 0.0083, Friedman; p = 0.0046, Wilcoxon), while the amplitude of 44-kHz calls remained stable (p = 0.0663, Friedman; p = 0.2661, Wilcoxon; 6^th^ ITI through 10^th^ ITI for both; Fig. 1S2E). After adjusting for angle-dependent hardware attenuation (see Methods, Sound mean power), the situation reversed (Fig. 1S2F). Both long 22-kHz and 44-kHz vocalizations showed similar amplitude levels during the first half of the fear conditioning session, while during the 6^th^-10^th^ ITI, 44-kHz calls were significantly louder than long 22-kHz calls (p = 0.0007-0.0097, Mann-Whitney).

### 44-kHz calls linked to freezing

We investigated the freezing behavior of all Wistar rats emitting 44-kHz vocalizations during 10 trials of fear conditioning (Tab. 1/Exp. 1-3/#2,4,8,13; n = 46). The training sessions were divided into 10-s-long time bins, from which we analyzed only the bins that had exclusively long 22-kHz or 44-kHz calls. For comparison, we also measured the freezing levels during the first 5 min of the trial (baseline freezing levels before any foot-shocks) as well as the bins in which animals did not vocalize (from the period after the 1^st^ shock to the end of the session). Of the n = 46 rats analyzed, n = 41 emitted 22-kHz vocalizations, from which n = 32 also emitted 44-kHz vocalizations, from which only n = 21 were determined to have both – 10-s-long bins of 22-kHz calls only and 44-kHz calls only (Tab. 2A). Freezing during the bins of 22-kHz calls only (p < 0.0001, for both groups) and during 44-kHz calls only bins (p = 0.0003) was higher than during the first 5 min baseline freezing levels of the session. Also, the freezing associated with emissions of 44-kHz calls only was higher than during bins with no ultrasonic vocalizations (p = 0.0353), and it was also 9.9 percentage points higher than during time bins with only long 22-kHz vocalizations, but the difference was not significant (p = 0.1907; all Wilcoxon).

To further investigate this potential difference, we measured freezing during the emission of randomly selected single 44-kHz and 22-kHz vocalizations. The minimal freezing behavior detection window was reduced to compensate for the higher resolution of the measurements (3, 5, 10, or 15 video frames were used). There was no difference in freezing during the emission of 44-kHz vs. 22-kHz vocalizations for ≥150-ms-long calls (3 frames, p = 0.2054) and for ≥500-ms-long calls (5 frames, p = 0.2404; 10 frames, p = 0.4498; 15 frames, p = 0.7776; all Wilcoxon, Tab. 2B).

### 44-kHz calls sorted into five subtypes

While the majority of 44-kHz vocalizations were of continuous unmodulated frequency (Fig. 2B), some comprised additional elements. Based on the composition of individual call elements and their relation to each other, we manually sorted the calls into five categories (Fig. 2B-F). If the start (*prefix*) or end (*suffix*) portion of a call was less than 1/5^th^ the length of the following or previous element, this portion of the call was not considered in its categorization into the five subtypes. The names and descriptions of the five subtypes are: **flat** – single element with near constant frequency and little to no interruptions to the sound continuity on the spectrogram; **step up** – two elements with an instantaneous frequency jump, where the first element is of lower frequency; **step down** – two elements with an instantaneous frequency jump, where the first element is of higher frequency; **insert** – three elements with an instantaneous frequency change, where the middle element is of different frequency; **complex** – more than three elements with instantaneous frequency changes.

### 44-kHz and 22-kHz calls closely related

44-kHz were emitted in aversive behavioral situations – as 22-kHz calls are observed (Antoniadis and McDonald, 1999, Dupin et al., 2019, Taylor et al., 2017). Both types of calls are long (usually >300 ms) and frequency-unmodulated. Some of the elements constituting such as step up; step down; insert and complex 44-kHz vocalizations (Fig. 2C-F) were at a lower frequency – typical for 22-kHz vocalizations. *Vice versa* we also observed 22-kHz calls with 44-kHz-like elements. Therefore, we propose that these long 22-kHz and 44-kHz vocalizations constitute a *supertype* group of long unmodulated aversive calls (“long 22/44-kHz vocalizations”).

We observed a stable, approximately 1.5 ratio in peak frequency levels between 22-kHz and 44-kHz vocalizations within individual rats. Specifically, in fourteen rats (13 Wistar and 1 SHR) with a clear transition from 22-kHz to 44-kHz calls during the fear conditioning session (n = 14, selected from Tab. 1/Exp. 1-3/#2,4,6-8,10-13), the proportion between the frequencies of the long 22-kHz vocalizations and the long 44-kHz calls was 1.48 ± 0.02. Similar results were obtained for 70 step up (1.53 ± 0.03) and 65 step down (1.59 ± 0.02) 44-kHz calls – altogether suggesting a 1.5-times or 3:2 frequency ratio. This ratio and its relevance has been observed in invertebrates and vertebrates including human speech and music (Hoeschele, 2017). In music theory, 3:2 frequency ratio is referred to as a perfect fifth and is often featured, e.g., the first two notes of the *Star Wars* 1977 movie (ascending, i.e., step up; comp. Fig. 2C, Track 1) and *Game of Thrones* 2011 television series (descending, i.e., step down; comp. Fig. 2D, Track 2) theme songs. All of which may point to a common basis for this sound interval and its prevalence which could be explained by the observation that all physical objects capable of producing tonal sounds generate harmonic vibrations, the most prominent being the octave, perfect fifth, and major third (Christensen, 1993, discussed in Bowling and Purves, 2015).

### 44-kHz calls separated in cluster analyses

Next, we showed that 44-kHz calls indeed constitute a separable type of ultrasonic vocalizations as it was sorted into isolated clusters by two different methods. First, using the DBSCAN algorithm method based on calls’ peak frequency and duration, we were able to divide all vocalizations recorded during all training sessions into 44-kHz vocalizations vs. all other vocalizations as two separate clusters (Fig. 3A). Secondly, a clustering algorithm that includes call contours, i.e., k-means with UMAP projection done via DeepSqueak (Figs 3BC, 3S1), sorted 44-kHz vocalizations of different subtypes including unusual ones (Fig. 2S1A-F), into topologically-separate groups. Notably, flat 44-kHz calls were consistently in a separate cluster from 22-kHz calls Figs 3C, 3S1B).

### 44-kHz calls formed separate groups

We have examined transition probabilities for call-type transitions within all vocalizations and within vocalization bouts in all Wistar rats during 10 trials of fear conditioning training (Tab. 1/Exp. 1-3/#2,4,8,13; n = 46). Transition probabilities between vocalization types were defined by counting the number of specific pair sequences and dividing by all observed sequence pairs where the first type of signal is followed by any signal. The most probable call following long 22-kHz vocalization was another long 22-kHz vocalization (94.4% probability). Similarly, the most probable call following 44-kHz vocalization was another 44-kHz call (83.1%). These values augmented when the analysis was limited to vocalizations emitted only in bouts, i.e., with <320 ms time gaps between calls (Wöhr et al., 2005), and reached 96.3% and 86.5%, respectively. Examples of parts of groups of both types of calls are demonstrated in Fig. 3S2.

### Response to 44-kHz playback

To describe the behavioral and physiological impact of 44-kHz vocalizations, we performed playback experiments in two separate groups of rats (Methods, Figs 4, 4S1). Overall, the responses to 44-kHz aversive calls presented from the speaker were either similar to 22-kHz vocalizations or in-between responses to 22-kHz and 50-kHz playbacks. For example, the heart rate of rats exposed to 22-kHz and 44-kHz vocalizations decreased, and increased to 50-kHz calls (Fig. 4A, comp. Olszyński et al., 2020). Whereas the number of vocalizations emitted by rats was highest during and after the playback of 50-kHz, intermediate to 44-kHz and lowest to 22-kHz playbacks (Figs 4BC, 4S1EF). Additionally, the duration of 50-kHz vocalizations emitted in response to 44-kHz playback was also intermediate, i.e., longer than following 22-kHz playback (Fig. 4D) and shorter than following 50-kHz playback (Figs 4D, 4S1G). Finally, similar tendencies were observed in the distance travelled and time spent in the half of the cage adjacent to the speaker (Fig. 4S1A-D).

**Fig. 4.**
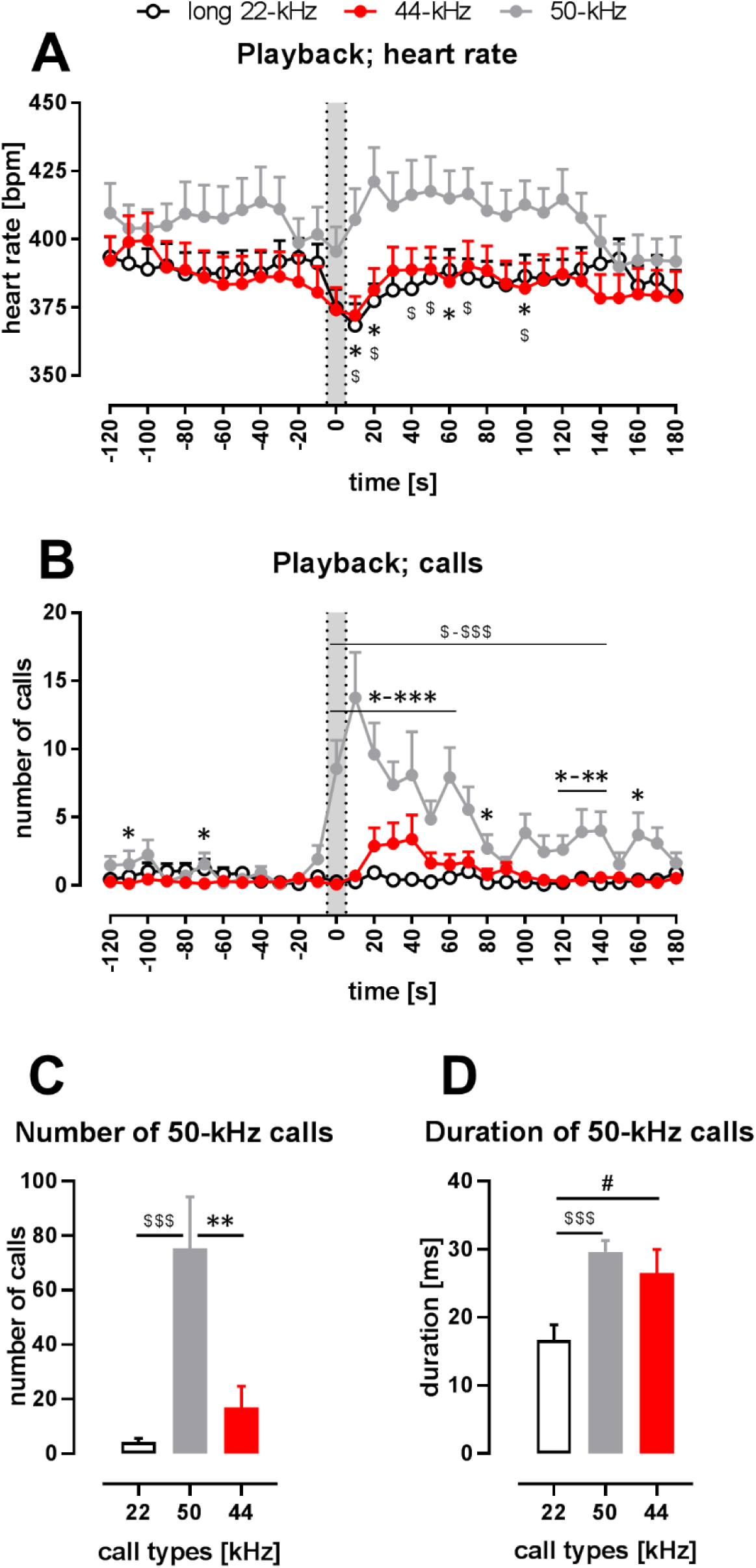
Physiological and behavioral response to playback of 44-kHz calls (vs. 50-kHz and 22-kHz calls) presented from a speaker to naïve Wistar rats. **A** – heart rate (HR); **B** – the number of emitted vocalizations. **AB** – gray sections correspond to the 10-s-long ultrasonic playback. Each point is a mean for a 10-s-long time-interval with SEM. **CD** – properties of 50-kHz vocalizations emitted in response to ultrasonic playback, i.e., number of calls (**C**) and duration (**D**) calculated from the 0-120 s range. **A** – 50-kHz playback resulted in HR increase (playback time-interval vs. 10-30 s time-interval, p = 0.0007), while the presentation of the aversive playbacks resulted in HR decrease, both in case of 22-kHz (p < 0.0001) and 44-kHz (p = 0.0014, average from -30 to -10 time-intervals (i.e., “*before*”) vs. playback interval, all Wilcoxon), which resulted in different HR values following different playbacks, especially at +10 s (p = 0.0097 for 50 kHz vs. 22-kHz playback; p = 0.0275 for 50 kHz vs. 44-kHz playback) and +20 s time-intervals (p = 0.0068, p = 0.0097, respectively, all Mann-Whitney). **B** – 50-kHz playback resulted also in a rise of evoked vocalizations (*before* vs. 10-30 s time-interval, p = 0.0002, Wilcoxon) as was the case with 44-kHz playback (p = 0.0176 in respective comparison), while no rise was observed following 22-kHz playback (p = 0.1777). However, since the increase in vocalization was robust in case of 50-kHz playback, the number of emitted vocalizations was higher than after 22-kHz playback (e.g., p < 0.0001 during 0-30 time-intervals) as well as after 44-kHz playback (e.g., p < 0.0001 during 0-10 time-intervals, both Mann-Whitney). Finally, when the increases in the number of emitted ultrasonic calls in comparison with *before* intervals were analyzed, there was a difference following 44-kHz vs. 22-kHz playbacks during 30 s and 40 s time intervals (p = 0.0420 and 0.0430, respectively, Wilcoxon). **C** – During the 2 min following the onset of the playbacks, rats emitted more ultrasonic calls during and after 50-kHz playback in comparison with 22-kHz (p < 0.0001) and 44-kHz (p = 0.0011) playbacks. The difference between the effects of 22-kHz and 44-kHz playbacks was not significant (p = 0.2725, comp. Fig. 4S1F; all Mann-Whitney). **D** – Ultrasonic 50-kHz calls emitted in response to playback differed in their duration, i.e., they were longer to 50-kHz (p = 0.0004) and 44-kHz (p = 0.0273, both Mann-Whitney) playbacks than to 22-kHz playback. * 50-kHz vs. 44-kHz, $ 50-kHz vs. 22-kHz, # 22-kHz vs. 44-kHz; one character (*, $ or #), p < 0.05; two, p < 0.01; three, p < 0.001; Mann-Whitney (**AB**) or Wilcoxon (**CD**). Values are means ± SEM, n = 13-16.

## Discussion

As Charles Darwin noted above (Darwin, 1872) and other researchers have confirmed (Briefer et al., 2012), the frequency level of animal calls is a vocal parameter that changes in accordance with its arousal state (intensity) or emotional valence (positive/negative state). The frequency shifts towards both higher and lower levels, i.e., alterations were observed during both positive (appetitive) and negative (agonistic/aversive) situations, however, as a general rule, frequency usually increases with an increase in arousal (Briefer et al., 2012). We would like to propose a hypothesis that our prolonged fear conditioning increased the arousal of the rats with no change in the valence of the aversive stimuli.

It could also be speculated that several other factors, apart from increased arousal, contributed to the emergence of 44-kHz vocalizations in our fear-conditioned rats, e.g., heightened fear, stress/anxiety, annoyance/anger, disgust/boredom, grief/sadness, despair/helplessness, and weariness/fatigue. It is not possible, at this stage, to definitively determine which factors played a decisive role. Please note that the potential contribution of these factors is not mutually exclusive.

However, several arguments support the idea that 44-kHz vocalizations communicate an increased negative emotional state. First, in general, ultrasonic vocalizations serve as a means of communicating rats’ emotional state (Brudzynski, 2013). Second, the changing of *the pitch of the voice bears some relation to certain states of feeling* (Darwin, 1872). Third, 44-kHz calls were notably more frequent during prolonged aversive stimulation, i.e., the 5^th^-10^th^ trials of fear conditioning. Fourth, they were linked to freezing. Fifth, they appeared as partial replacements of, established as aversive, 22-kHz calls – in the presence of the same painful stimulus. Sixth, numerous instances of vocalizations featured both 22-kHz-like and 44-kHz-like call-elements.

Also, several observations contradict the potential contribution of fatigue. The sound mean power of 44-kHz vocalizations was comparable to, or possibly even higher than, that of 22-kHz calls, despite the higher energy costs associated with producing higher-pitched calls (Sonninen and Hurme, 1998), i.e., the rats emitting 44-kHz calls invested additional energy to communicate their emotional state; both *in vivo* measurements (Riede, 2013) and computer modelling (Hakansson et al., 2022) demonstrated that producing calls of higher frequency, such as 50 kHz vs. 22 kHz, requires increased activity of various muscles. Additionally, the mean power of 44-kHz vocalizations remained strong and stable for several trials – in contrast to 22-kHz vocalizations. Finally, when 44-kHz calls started to appear in significant numbers, i.e., after the 4^th^-5^th^ trials of fear conditioning, they were as long as 22-kHz vocalizations.

Concerning the latter, we observed a significant decrease in the mean power of 22-kHz vocalizations during the fear conditioning session. Such reduction could potentially be attributed to fatigue (as observed in humans, Kitch and Oates, 1994), despair (e.g., as a reaction to the lack of effects from repeated emissions of 22-kHz calls), or both. The reduction in the amplitude of 22-kHz calls during the 10-trial fear conditioning was also recently observed by others (Gonzalez-Palomares et al., 2023).

Importantly, it has been demonstrated multiple times that training rats with several electric foot-shocks (i.e., 5-10 shocks) produces a qualitatively different kind of fear-memory compared to training with only 1-2 shocks. Training with more numerous shocks has been shown to result in augmented freezing (e.g., Fanselow and Bolles, 1979, Haubrich et al., 2020, Haubrich and Nader, 2023, Poulos et al., 2016, Wang et al., 2009) which reflects a more intense fear-memory that is resistant to extinction (Haubrich et al., 2020, Haubrich and Nader, 2023), resistant to reconsolidation blockade (Haubrich et al., 2020, Wang et al., 2009, Finnie and Nader, 2020), associated with downregulation of NR2B NMDA-receptor subunits as well as elevated amyloid-beta concentrations in the lateral and basal amygdala (Finnie and Nader, 2020, Wang et al., 2009). Additionally, it involves activation of the noradrenaline-locus coeruleus system (Haubrich et al., 2020) and collective changes in connectivity across multiple brain regions within the neural network (Haubrich and Nader, 2023).

Notably, it has also been shown that higher freezing as a result of fear-conditioning training correlates with increased concentrations of stress hormone, corticosterone, in the blood (Dos Santos Correa et al., 2019). The rats subjected to 6- and 10-trial fear conditioning, whose results are reported herein (Tab. 1/Exp. 2/#7,8,11,12; n = 73), also demonstrated higher freezing than rats subjected to 1-trial conditioning (Tab. 1/Exp. 2/#6,10; n = 33), which is reported elsewhere (Olszynski et al., 2021, Fig. S1C-E; Olszynski et al., 2022, Fig. S1D-G). Therefore, we postulate that emission of 44-kHz calls is associated with increased stress and the training regime forming robust memories.

Amounting research points to the utility of rat ultrasonic vocalizations to alter emotional states, evidenced by behavioral changes, in tested rats *via* playback of affectively valenced calls (Bonauto et al., 2023). We have exposed rats to 44-kHz playback along with 22-kHz and 50-kHz playback. The experimental design (see methods for details) allowed us to compare rats’ responses to 22-kHz vs. 44-kHz playbacks especially – with 50-kHz playback used as a form of control or baseline. In general, the rats responded similarly to hearing 44-kHz calls as they did to hearing aversive 22-kHz calls, especially regarding heart-rate change, despite the 44-kHz calls occupying the frequency band of appetitive 50-kHz vocalizations. This is contrary to some observations (Saito et al., 2019) which suggested that frequency band plays the main role in rat ultrasound perception. Please factor in potential carry-over effects (resulting from hearing playbacks of the same valence in a row) in the differences between responses to 50-kHz vs. 22/44-kHz playbacks, especially, those observed before the signal (Fig. 4AB). Other responses to 44-kHz calls were intermediate, they fell between response levels to appetitive vs. aversive playback, which might signify some behavioral specificity and importance (or possibly confusion). These latter effects were similar in both playback experiments despite an array of methodological differences between them. Overall, these initial results raise further questions about how, ethologically, animals may interpret the variation in hearing 22-kHz vs. 44-kHz calls and integrate this interpretation in their responses.

The question also is, why have the 44-kHz vocalizations been overlooked until now? On one hand, long (or not that long as in Biały et al., 2019), frequency-stable high-pitch vocalizations have been reported before (e.g., Sales, 1979; Shimoju et al., 2020), notably as caused by intense cholinergic stimulation (Brudzynski and Bihari, 1990) or higher shock-dose fear conditioning (Wöhr et al., 2005). However, they have not been systematically defined, described, fully shown or demonstrated to be a separate type of vocalization. On the other hand, 44-kHz calls were likely omitted as the analyses were restricted to canonical groups, i.e. flat 22-kHz and short 50-kHz calls, with a sharp dividing frequency border between the two (e.g., Kalamari et al., 2021, Potasiewicz et al., 2020, Turner et al., 2019) or even a frequency ‘safety gap’ between 22-kHz and 50-kHz vocalizations (e.g., Silkstone and Brudzynski, 2019, Garcia et al., 2015). Moreover – many older bat-detectors had limited frequency-range detection (e.g., up to 40 kHz in Sales, 1991), when stress-evoked types of ultrasonic calls were being established. Finally, 44-kHz vocalizations are emitted much fewer than 22-kHz calls (Fig. 1FG).

Here we present introductory evidence that 44-kHz vocalizations are a separate and behaviorally-relevant group of rat ultrasonic calls. These results require further confirmations and additional experiments, also in form of replication, including research on female rat subjects. However, our results bring to awareness that rats employ these previously unrecognized, long, high-pitched and flat aversive calls in their vocal repertoire. Researchers investigating rat ultrasonic vocalizations should be aware of their potential presence and to not rely fully on automated detection of high vs. low-pitch calls.

## Materials and Methods

### Animals

Wistar rats (n = 167) were obtained from The Center for Experimental Medicine of the Medical University of Bialystok, Poland; spontaneously hypertensive rats (SHR, n = 80) and Sprague-Dawley rats (n = 16) were from Mossakowski Medical Research Institute, Polish Academy of Sciences, Poland. All rats were males, 7 weeks of age on arrival, randomly assigned into groups and cage pairs where appropriate; housed with a 12 h light-dark cycle, ambient temperature (22–25 °C) with standard chow and water provided *ad libitum.* The animals were left undisturbed for at least one week before any procedures, then handled at least four times for 2 min by each experimenter directly involved for one to two weeks. All procedures were approved by Local Ethical Committees for Animal Experimentation in Warsaw.

### Animal details: groups of animals used

#### Trace fear conditioning experiment

Wistar rats, both single-housed (n = 14) and pair-housed (n = 20), were implanted with radiotelemetric transmitters for measuring heart rate in an ultrasonic vocalization playback experiment previously described by us (Olszyński et al., 2020) after which, at 13 weeks of age, half of them (n = 17) were fear-conditioned (10 shocks), while the other half (n = 17) served as controls (Tab. 1/Exp. 1/#1-4, n = 34).

#### Delay fear conditioning experiment, rats with transmitters

Wistar rats (n = 94) and SHR (n = 80) were implanted with a radiotelemetric transmitters one week before fear conditioning during which they received 0, 1, 6 or 10 shocks at 12 weeks of age (Tab. 1/Exp. 2/#5-12, n = 174). All the details are described in Olszyński et al. (2021) and Olszyński et al. (2022).

#### Delay fear conditioning experiment, rats without transmitters

Wistar rats were housed in pairs; were not implanted with radiotelemetric transmitters to eliminate the potential effect of surgical intervention on vocalization; they received 10 conditioning stimuli at 12 weeks of age (Tab. 1/Exp. 3/#13, n = 10) – same as in Olszyński et al. (2021) and Olszyński et al. (2022).

#### Playback experiment, rats with transmitters

Wistar rats (n = 29) were housed in pairs; all were implanted with a radiotelemetric transmitter one week before the playback experiment. At 12 weeks of age, one group (n = 13) heard 50-kHz appetitive vocalization playback while the other (n = 16) 22-kHz and 44-kHz aversive calls (for details see below).

#### Playback experiment, rats without transmitters

Sprague Dawley rats (n = 16) were housed in pairs, were not implanted with the transmitters, and received 22-kHz, 44-kHz, and 50-kHz ultrasonic vocalization playback at 8 weeks of age (see below).

### Surgery, transmitter implantation, heart-rate registration

Radiotelemetric transmitters (HD-S10, Data Sciences International, St. Paul, MN, USA) were implanted into the abdominal aorta of rats in specified groups as previously described (Olszyński et al., 2020, Olszyński et al., 2021). An illustrative image with the surgery details can be found elsewhere (Figure 5 in Pestana-Oliveira et al., 2020; please note, tissue glue was used instead of cellulose patches and silk sutures). The signal was collected by receivers (RSC-1, Data Sciences International, St. Paul, MN, USA) as previously described (Olszyński et al., 2020, Olszyński et al., 2021, Olszyński et al., 2022). Readings were processed using Dataquest ART (version 4.36, Data Sciences International) for trace fear conditioning (Tab. 1/Exp. 1) and Ponemah (version 6.32, Data Sciences International) software for other experiments (Tab. 1/Exp. 2-3 and playback experiments).

### Fear conditioning

All conditioning procedures were conducted in a chamber (VFC-008-LP, Med Associates, Fairfax, VT, USA) located in an outer cubicle (MED-VFC2-USB-R, Med Associates) equipped with an ultrasound CM16/CMPA condenser microphone (Avisoft Bioacoustics, Berlin, Germany). Ultrasonic vocalizations were recorded via Avisoft USGH Recorder (Avisoft Bioacoustics), and rat behavior was recorded via NIR monochrome camera (VID-CAM-MONO-6, Med Associates). All procedures were described in detail before (Olszyński et al., 2021, Olszyński et al., 2022).

#### Trace fear conditioning

(Tab. 1/Exp. 1/#1-4, n = 34 rats) was performed similarly to some previous reports (e.g., Jahołkowski et al., 2009). Rats were individually placed in the fear conditioning apparatus in one of two different contexts: A (safe) or B (unsafe). Context A was in an illuminated room with the cage interior with white light, the cage floor was made of solid plastic, and the cage was scented with lemon odor, cleaned with a 10% ethanol solution; the experimenter was male wearing white gloves. Context B was a different, dark room, with the cage interior with green light, the floor was made of metal bars, and the cage was scented with mint odor, cleaned with 1% acetic acid; the experimenter was female with violet gloves. The procedure: on day -2, each rat was habituated to context A for 20 min; on day -1, habituated to context B for 20 min; on day 0, each rat was placed for 52 min in context A; on day 1, after 10 min in context B, the rat received 10 conditioning stimuli (15-s-long sine wave tone, 5 kHz, 85 dB) followed by a 30 s trace period and a foot-shock (1 s, 1 mA) and 210 s inter-trial interval, i.e., ITI; total session duration: 52 min. Control rats were subjected to the same procedures but did not receive the electric shock at the end of trace periods. The animals were tested with the same protocol without shocks in context A (day 2) and context B (day 3). During the test session, control animals showed a lower level of freezing than conditioned animals (1.3 ± 0.8% vs. 19.7 ± 4.3% during the first 5 min in unsafe context B and 0.4 ± 0.3% vs. 9.9 ± 1.9% during 10 s following the time of expected shock in context B, results averaged from the first 3 out of 10 trials; p = 0.0003 and p = 0.0001, respectively, Mann-Whitney); none of the control animals emitted 44-kHz calls, neither the fear conditioning day nor the test days.

#### Delay fear conditioning

(Tab. 1/Exp. 2-3/#5-13, n = 184 rats) The procedure and its results were described before (Olszyński et al., 2021, Olszyński et al., 2022); rats received 1, 6 or 10 conditioning stimuli (20-s-long white light co-terminating with an electric foot-shock, 1 s, 1 mA). For control rats, an equal time-length procedure was done for each conditioning protocol, i.e., the same parameters as in 1, 6 or 10 stimuli groups, with no shock. Control animals showed a lower level of freezing than conditioned animals. There were only 4 ultrasonic calls we classified as 44-kHz vocalizations among 4,126 vocalizations emitted by the control rats during training and testing. We did not observe any difference in the number of 44-kHz vocalizations between Wistar rats with transmitters vs. without transmitters during delay conditioning training (p = 0.8642, Mann-Whitney). These two groups were therefore reported together.

### Measuring freezing

Freezing behavior was scored automatically using Video Freeze software (Med Associates) with a default motion index threshold of 18. To avoid including brief moments of the animal’s stillness, freezing was measured only if the animal did not move for at least 1 s, i.e., 30 video frames, with some exceptions, see next.

#### Vocalization-nested freezing behavior

Freezing at the exact times of ultrasonic calling was measured in rats that had undergone 10 trials of fear conditioning which produced 44-kHz calls (n = 32, selected from Tab. 1/Exp. 1-3/#2,4,8,13). From each rat, one 44-kHz call was randomly selected along with the long 22-kHz call closest to it. Such pairs of vocalizations were selected with either ≥150 ms duration (n = 32) or ≥500 ms duration (n = 28). For each pair of vocalizations, the freezing behavior was calculated from the entire duration of the shorter call and for the equal-time-length period in the middle of the longer vocalization. Due to the shortened time-scale, the minimal freezing detection window was reduced to 3 frames for ≥150-ms-long calls as well as 5, 10, and 15 frames – for ≥500-ms calls.

### Ultrasonic playback

It was performed as described previously (Olszyński et al., 2020, Olszyński et al., 2021, Olszyński et al., 2022) in individual experimental cages with acoustic stimuli presented through a Vifa ultrasonic speaker (Avisoft Bioacoustics, Berlin, Germany) connected to an UltraSoundGate Player 116 (Avisoft Bioacoustics). Ultrasonic vocalizations emitted by the rat were recorded by a CM16/CMPA condenser microphone (Avisoft Bioacoustics). Both playback and recording of calls were performed using Avisoft Recorder USGH software (version 4.2.28, Avisoft Bioacoustics). The locomotor activity was recorded with an acA1300-60gc camera (Basler AG, Ahrensburg, Germany). There were 8 sets of ultrasonic vocalizations presented:

1. **44-kHz long calls**, 8 calls in 1 repeat, constant frequency (2.7 ± 0.1 kHz max-min frequency difference), 42.1 ± 0.2 kHz peak frequency, 1064.3 ± 89.6 ms duration with 199.0 ± 14.7 ms sound intervals;
2. **22-kHz long calls**, 8 calls in 1 repeat, typical long 22-kHz vocalizations, constant frequency (1.9 ± 0.9 kHz max-min frequency difference), 24.5 ± 0.2 kHz peak frequency, 1066.4 ± 90.2 ms duration with 195.6 ± 15.5 ms sound intervals;
3. **22-kHz short modulated calls**, 26 calls in 2 repeats, short (<300 ms), not resembling typical 22-kHz long calls (5.3 ± 0.4 kHz max-min frequency difference), 22.7 ± 0.6 kHz peak frequency, 24.7 ± 1.6 ms duration with 172.8 ± 5.6 ms sound intervals;
4. **22-kHz short flat calls**, 43 calls in 1 repeat, short (<300 ms), resembling typical 22-kHz long calls, constant frequency (2.3 ± 0.1 kHz max-min frequency difference), 25.1 ± 0.3 kHz peak frequency, 102.4 ± 10.9 ms duration with 132.1 ± 6.2 ms sound intervals;
5. **50-kHz modulated calls**, 23 calls in 2 repeats, moderately modulated (8.6 ± 0.3 kHz max-min frequency difference), 61.0 ± 0.8 kHz peak frequency, 37.6 ± 1.5 ms duration with 183.7 ± 4.5 ms sound intervals;
6. **50-kHz flat calls**, 29 calls in 2 repeats, constant frequency (4.2 ± 0.2 kHz max-min frequency difference), 53.5 ± 0.5 kHz peak frequency, 66.2 ± 3.8 ms duration with 144.1 ± 4.4 ms sound intervals;
7. **50-kHz trill calls**, 29 calls in 2 repeats, highly modulated (37.4 ± 1.7 kHz max-min frequency difference), 68.0 ± 0.9 kHz peak frequency, 53.7 ± 1.4 ms duration with 158.5 ± 4.9 ms sound intervals;
8. **50-kHz kHz mixed calls**, used previously in Olszyński et al. (2020), Olszyński et al. (2021), and Olszyński et al. (2022), 28 calls, in 3 repeats, frequency modulated and trill subtypes, 9.8 ± 1.9 kHz max-min frequency difference, 58.6 ± 0.7 kHz peak frequency, 28.4 ± 1.6 ms duration with 91.4 ± 1.4 ms sound intervals.

Calls were presented with a sampling rate of 250 kHz in 16-bit format. All calls except for 50-kHz mixed calls were collected in our laboratory from fear conditioning or playback experiments. Calls in the same set were taken from one animal wherever possible. The sound interval was adjusted if it was peculiarly long or the sequence was interrupted by other types of calls in the original recordings.

**Playback procedure, rats with transmitters;** as previously described (Olszyński et al., 2020, Olszyński et al., 2021, Olszyński et al., 2022). Before playback presentation, animals were habituated for 3 min to the experimental conditions, i.e., recording cage, presence of the speaker and microphone, over 4 days. Habituated rats then underwent a playback procedure, in short, after 10 min of silence, the rats were exposed to four 10-s-long call sets (either aversive or appetitive) with 5-min-long ITI in-between; a rat that received appetitive playback was followed by a rat receiving aversive playbacks etc. Also, the order of the presented sets was randomized between animals. The aversive-calls playback contained sets nos. 1-4. The appetitive-calls playback contained sets nos. 5-8. Since initial analysis showed no differences within responses to 22-kHz aversive sets and within responses to 50-kHz appetitive sets, we decided to show the results following playback of 44-kHz long calls (set no. 1), 22-kHz long calls (set no. 2), and 50-kHz modulated calls (set no. 5) only.

#### Playback procedure, rats without transmitters

Before playback presentation, animals were habituated for 3 min to the experimental conditions, i.e., recording cage, presence of the speaker and microphone, over 4 days. After 5 min of initial silence, the rats were presented with two 10-s-long playback sets of either 22-kHz (set no. 2; n = 8) or 44-kHz calls (set no. 1; n = 8), followed by one 50-kHz modulated call 10-s set (no. 5) and another two playback sets of either 44-kHz or 22-kHz calls not previously heard. The playback presentations were separated by 3 min ITI. Responses to the pairs of playback sets were averaged.

#### Locomotor activity in playback

An automated video tracking system (Ethovision XT 10, Noldus, Wageningen, The Netherlands) was used to measure the total distance travelled (cm). Proximity to the speaker was expressed as the percentage of time spent in the half of the cage closer to the ultrasonic speaker. Center-point of each animal’s shape was used as a reference point for measurements of locomotor activity thus registering only full-body movements.

### Analysis of ultrasonic vocalizations

Audio recordings were analyzed manually using SASLab Pro (version 5.2.xx, Avisoft Bioacoustics) as described (Olszyński et al., 2020, Olszyński et al., 2021, Olszyński et al., 2022) to measure key features of calls and categorize them into subtypes.

**Sound mean power** was measured as the average spectra power density of the vocalization contour using DeepSqueak software. Initially, calls were detected using the default rat long-vocalization neural network (Long Rat Detector YOLO R1) and subsequently manually reviewed and corrected where necessary. We analyzed a subset of Wistar rats subjected to 10-trial fear conditioning that emitted more than 20 instances of 44-kHz calls during the fear conditioning session (n = 17, selected from Tab. 1/Exp. 1-3/#2,4,8,13). It is important to note that due to the directional characteristics of the microphones used, angular attenuation occurred during audio recording. This phenomenon results in a selective reduction in the intensity of higher frequency sounds, dependent on the angle between the sound emitter and the microphone (as specified in the CM16/CMPA microphone hardware specification page, Avisoft Bioacoustics website). In our experimental setup, we approximated a 45° angle between the plane of the rat’s head and the plane of the microphone’s membrane. This angle corresponds to an estimated 10 dB attenuation (adopting a conservative estimate) of 40-kHz frequencies compared to 20-kHz frequencies for which there is even a small dB gain due to these hardware properties, 44-kHz calls are predicted to be approximately at least 10 dB louder in reality than what was recorded.

### 22-kHz vs. 44-kHz frequency ratio

A clear transition point between 22-kHz and 44-kHz long calls was observed in n = 13 Wistar rats and n = 1 SHR. In each case, ten 22-kHz calls followed by ten 44-kHz calls were analyzed (n = 14, selected from Tab. 1/Exp. 1-3/#2,4,6-8,10-13).

#### Step up and step down frequency ratio

Rats which emitted at least five vocalizations of the specific subtype were analyzed (step up, n = 14; step down, n = 13; selected from Tab. 1/Exp. 1-3/#2,4,7,8,13; 5 calls of the two subtypes from each rat were chosen randomly and the frequencies of their elements were measured.

**Ultrasonic vocalizations clustering** (two independent methods)

Calls of conditioned and control animals were taken from all fear conditioning training sessions (Tab. 1/Exp. 1-3, n = 218). We used **DBSCAN algorithm** (Ester et al., 1996); a density based method, from the scikit-learn (sklearn) Python package, because of its ability to detect a desired number of clusters of arbitrary shape; with two main input parameters: MinPts (minimal number of points forming the core of the cluster) and ε (the maximum distance two points can be from one another while still belonging to the same cluster). To avoid detecting small clusters, we limited MinPts to 150 samples. The heuristic method described by Ester et al. (Ester et al., 1996) was implemented to find the initial range of *ε*. All the input data were standardized. The silhouette coefficient (Rousseeuw, 1987) was used to control the quality of the clustering. Maximizing *ε* among different ranges helped to select the most relevant number of identified clusters. Clustering with *ε* in the range of 0.14–0.2 resulted in a silhouette coefficient around 0.2–0.5.

#### K-means algorithm

Vocalizations of selected fear-conditioned rats with 6-10 shocks and >30 of 44-kHz calls (n = 26, selected from Tab. 1/Exp. 1-3/#2,4,7,8,11-13) were detected using a built-in neural network for long rat calls (Long Rat Detector YOLO R1) on DeepSqueak (Coffey et al., 2019) software (version 3.0.4) running under MATLAB (version 2021b, MathWorks, Natick, MA, USA) and manually revised for missed and mismatched calls. Unsupervised k-means clustering was based on call contour, frequency and duration variables, with equal weights assigned, and several descending elbow optimization parameters were used to obtain different maximum numbers of clusters together with Uniform Manifold Approximation and Projection for Dimension Reduction (UMAP) (McInnes et al., 2018) for superimposing and visualization of clusters.

### Quantification and statistical analysis

Data were analyzed using non-parametric Friedman, Wilcoxon, Mann-Whitney tests with GraphPad Prism 8.4.3 (GraphPad Software, San Diego, CA, USA); the p values are given, p *<* 0.05 as the minimal level of significance. In particular, the Friedman test was used to assess the presence of change within the sequence of several ITI, while the Wilcoxon test was used for the difference between the first and the last ITI analyzed. Figures were prepared using the same software and depict average values with a standard error of the mean (SEM).

## Data availability

Raw data (calls’ peak frequency and duration) analyzed, ultrasonic playback files used (.wav), data supporting clustering files for DBSCAN (.csv), and extracted call contours for k-means (.mat) have been deposited to Mendeley Data at http:/to be provided/. The other data in this study are available from the corresponding author upon request.

## Acknowledgements

We thank Iryna Artemieva for her help with DeepSqueak analysis. We would also like to thank Patrick Reilly and Adelaide Yiu for their advice and assistance. This research was funded by the National Science Centre, Poland, grant OPUS no. 2015/19/B/NZ4/03393 (R.K.F.) and by Mossakowski Medical Research Institute, PAS, Poland, Internal Research Fund no. FBW-17 (R.K.F.). R.P and A.D.W were supported by ESF, POWR.03.02.00-00-I028/17-00.

## Author contributions

K.H.O, and R.P., and R.K.F. designed the study and wrote the manuscript. K.H.O., R.P., A.D.W., A.W.G., and O.G. performed the experiments. W.P. and M.K. performed DBSCAN analysis. R.P. performed k-means analysis. K.H.O., R.P., I.A.Ł., and A.D.W. analyzed the data. R.K.F. acquired the funding and supervised the project. All authors reviewed and approved the final version of the manuscript.

^1^*peak frequency* of a given vocalization is defined here as the highest power peak in the averaged spectrum of the entire element.

---

**Fig. 1S1.**
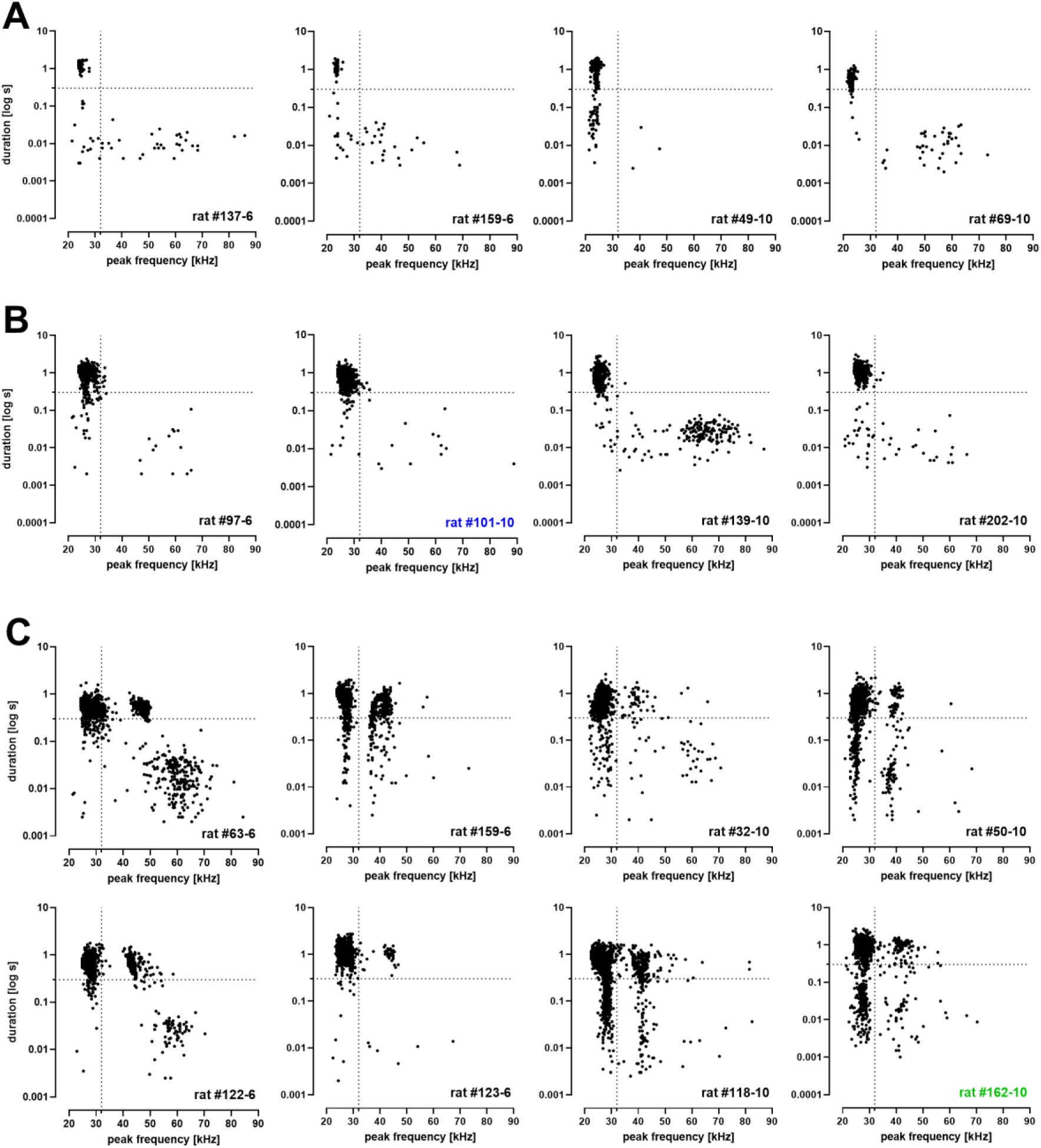
Variations of call frequency; shown in relation to call duration in Wistar rats that undergone 6 or 10 trials of delay fear conditioning. (n = 16, selected from Tab. 1/Exp. 2-3/#7,8,13). Vocalizations plotted in relation to peak frequency (x axis) and duration (y axis). Each point corresponds to one vocalization. Vertical dotted line marks threshold value (32 kHz) between 22-kHz and 50-kHz calls. Horizontal dotted line marks threshold value (300 ms) between short and long 22-kHz calls (Brudzynski et al., 1993). Rat identifier is given in lower right corner; the number after dash indicates the number of conditioning trials. **A** – examples from four rats which emitted typical long 22-kHz calls (no 44 kHz calls). **B** – four typical long 22-kHz vocalizations with few long 22-kHz calls crossing the 32 kHz threshold. **C** – eight sample rats which emitted typical long 22-kHz vocalizations and atypical high-frequency aversive calls forming a separate 44-kHz group.

**Fig. 1S2.**
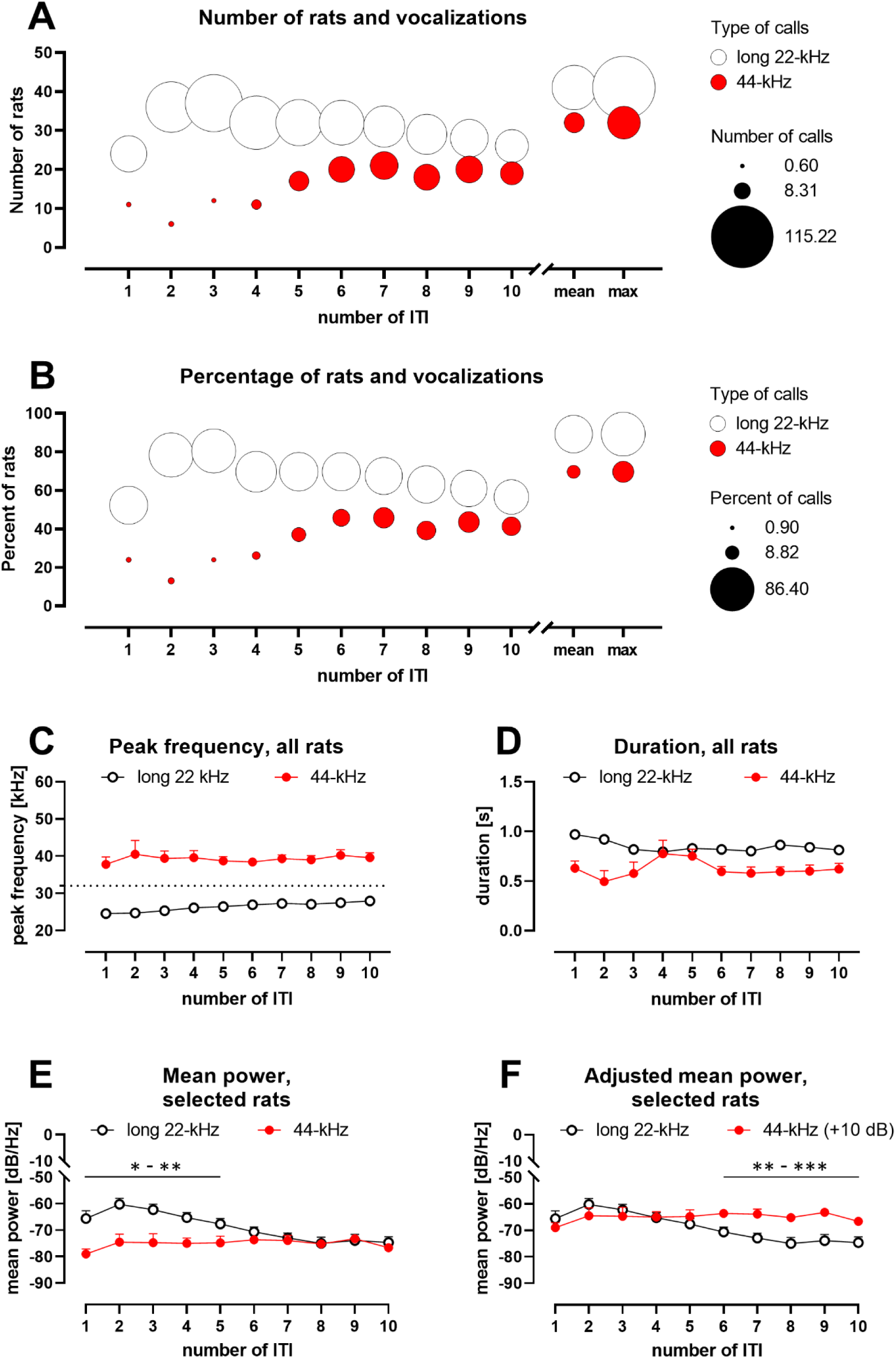
Changes in distribution (AB), frequency (C), duration (D), and mean power (EF) of long aversive vocalizations throughout fear conditioning session. Data were acquired from all Wistar rats subjected to a 10-trial fear conditioning procedure (Tab. 1/Exp. 1-3/#2,4,8,13; n = 46). X-axes represent subsequent inter-trial intervals (ITI) numbered after the preceding conditioned stimulus. **AB**. **Number or percentage of rats emitting long vocalizations.** Bubbles represent long 22-kHz calls (white) or 44-kHz calls (red); bubble size scales with the amount of vocalizations. Emission of 44-kHz calls and the number of animals emitting them increases in the latter half of the session. Data are absolute values (**A**) or percentages (**B**); “mean”, average value from all ITI, “max”, maximum values from each rat. **C**. **Frequencies of 22-kHz and 44-kHz vocalizations.** Horizontal dotted line marks the threshold value (32 kHz) between 22-kHz and 50-kHz/44-kHz calls. Peak frequency of long vocalizations rose gradually in all rats. **D. Duration of 22-kHz and 44-kHz vocalizations.** The duration of 22-kHz calls gradually declined. The duration of 44-kHz calls peaked after the 4^th^ ITI. **E. Mean power of 22-kHz and 44-kHz vocalizations.** Mean power spectral density (loudness, amplitude) of 22-kHz calls, n = 14-17 per ITI; and 44-kHz calls, n = 5-17 per ITI; **F.** results for 44-kHz vocalizations (from **E**) were adjusted for angular attenuation, i.e., +10 dB. Before the adjustment: during the first half of the session, 22-kHz calls appeared louder than 44-kHz calls, in the second half of the session the difference dissipated. After the adjustment: both types of calls started on a comparable amplitude level, but in the 6^th^-10^th^ ITI, 22-kHz calls became quieter than 44-kHz calls. Values are means ± SEM (**C-F**). Graphs show either all rats (**A-D**, n = 46) or rats which met the criteria of emitting >20 of 44-kHz calls (**EF**, n = 17 selected from n = 46); *p < 0.05, **p < 0.01, ***p < 0.001).

**Fig. 1S3.**
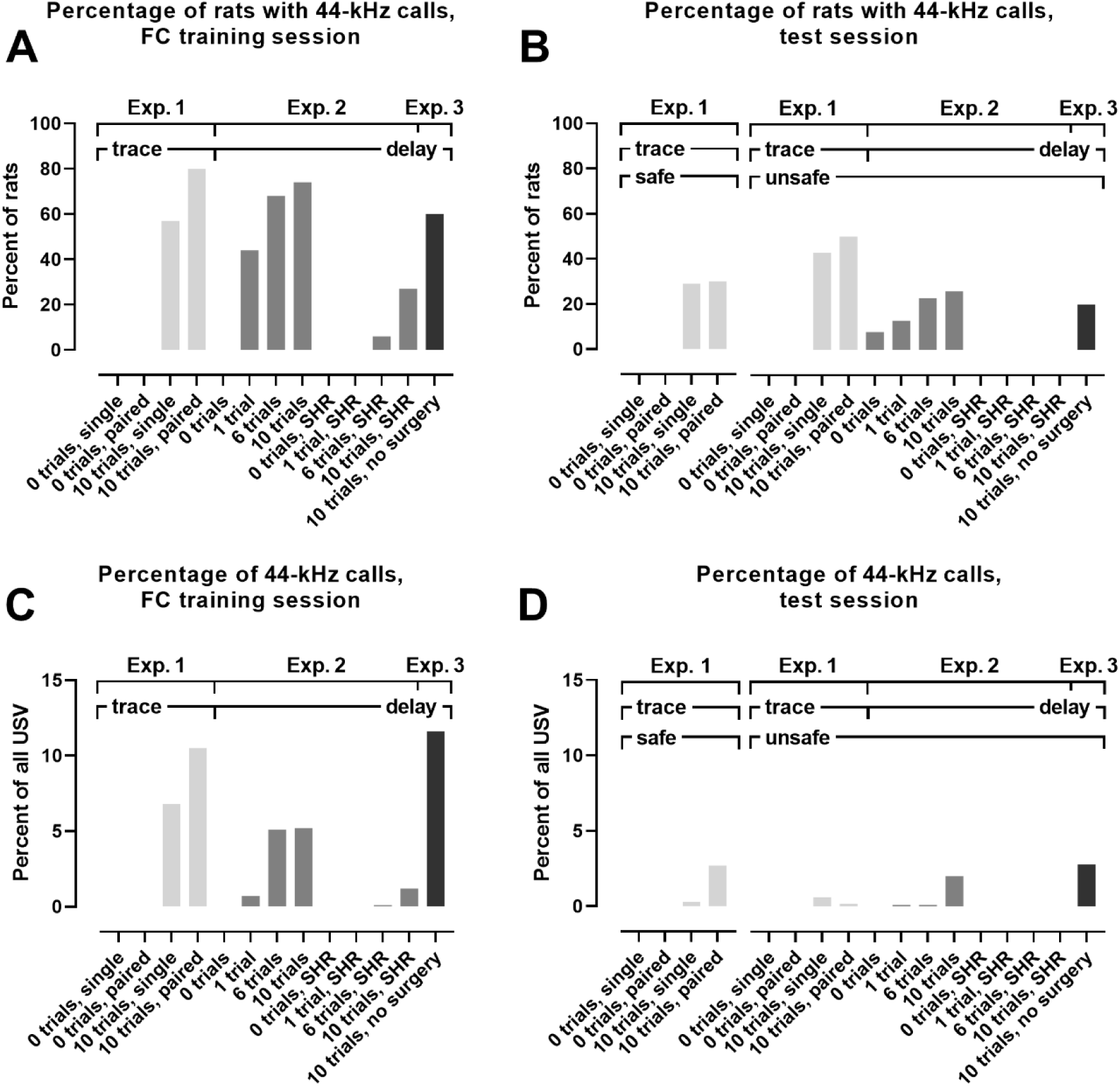
Percentage of animals emitting 44-kHz calls (AB) and percentage of 44-kHz calls in all vocalizations (CD) emitted by Wistar rats and SHR. Results from three main fear conditioning experiments are shown (comp. Tab. 1), i.e., Exp. 1 (light gray bars), Exp. 2 (dark gray bars), and Exp. 3 (black bars), which all were performed with Wistar rats or SHR (when specified in the x-axis labels). The labels denote different experimental groups used across the experiments (see Tab. 1 for the number of animals in each group). Results were obtained during fear conditioning training (**A**, **C**) and testing sessions (**B**, **D**). Rats subjected to trace fear conditioning were tested in safe and unsafe contexts, while in delay fear conditioning, the rats were tested only in an unsafe context (see Methods). 44-kHz calls appeared most often in Wistar rats which had undergone 10-trial fear conditioning procedures. Please note that the experiments were not performed in parallel.

**Fig. 2S1.**
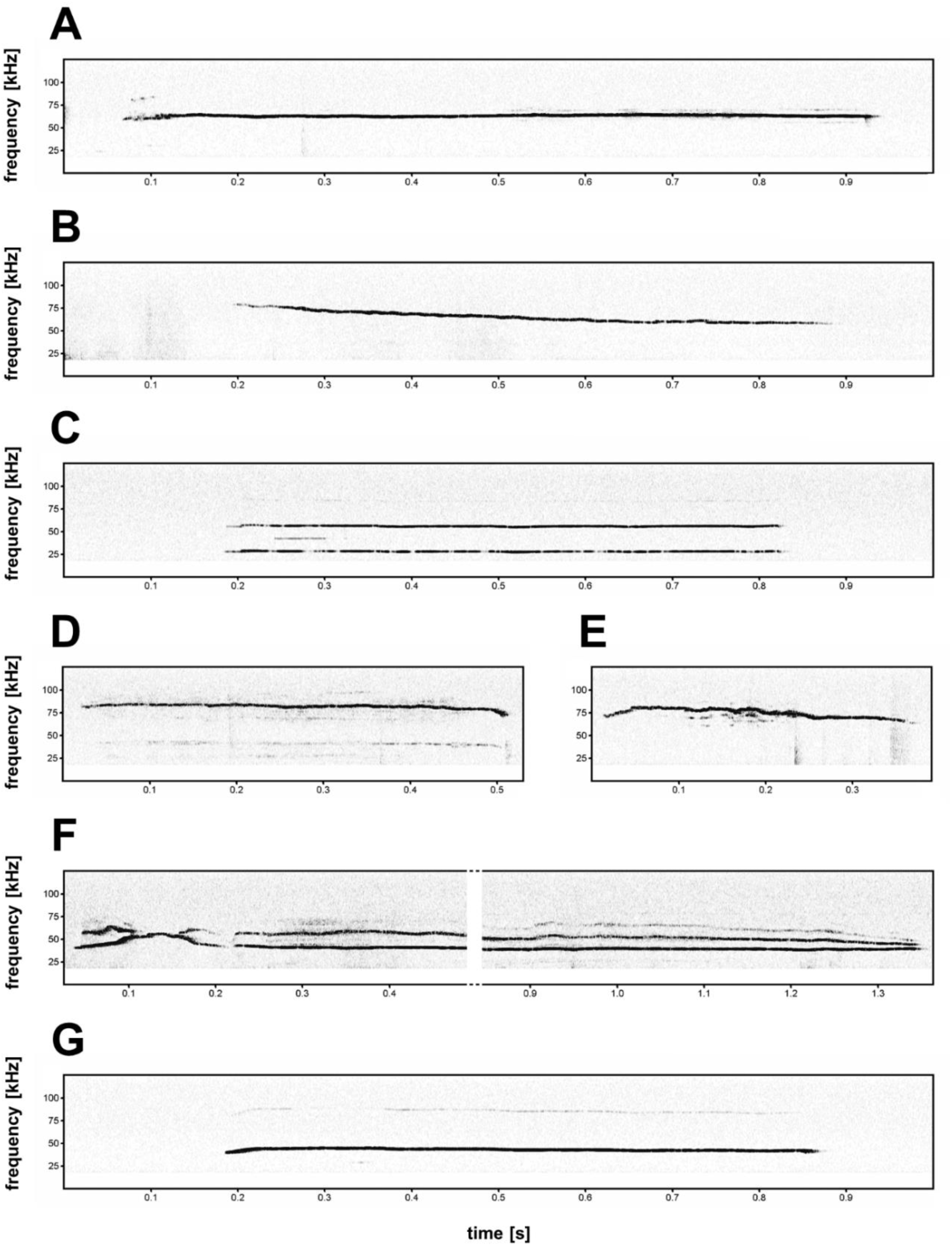
Non-typical 44-kHz aversive vocalizations. **A**, **B** – constant frequency calls with very high peak frequency (**A**, peak frequency = 62.9 kHz; **B**, peak frequency = 65.9 kHz, start peak frequency = 78.1 kHz). **C**, **D** – harmonic aversive vocalizations, where element with fundamental frequency (F0, lowest frequency of the vocalization) is not with maximum amplitude, i.e., peak frequency is determined from the higher call component (**C**, F0 = 27.8 kHz, peak frequency = 55.6 kHz; **D**, F0 = 40 kHz, peak frequency = 81.5 kHz). **E**, **F** – vocalizations with prominent duration but with modulated frequency (**E**, peak frequency = 69.3 kHz; **F**, peak frequency = 39.0 kHz). **A**, **G** – constant frequency calls from SHR (**G**, flat 44-kHz call, peak frequency = 42.4 kHz).

**Fig. 3S1.**
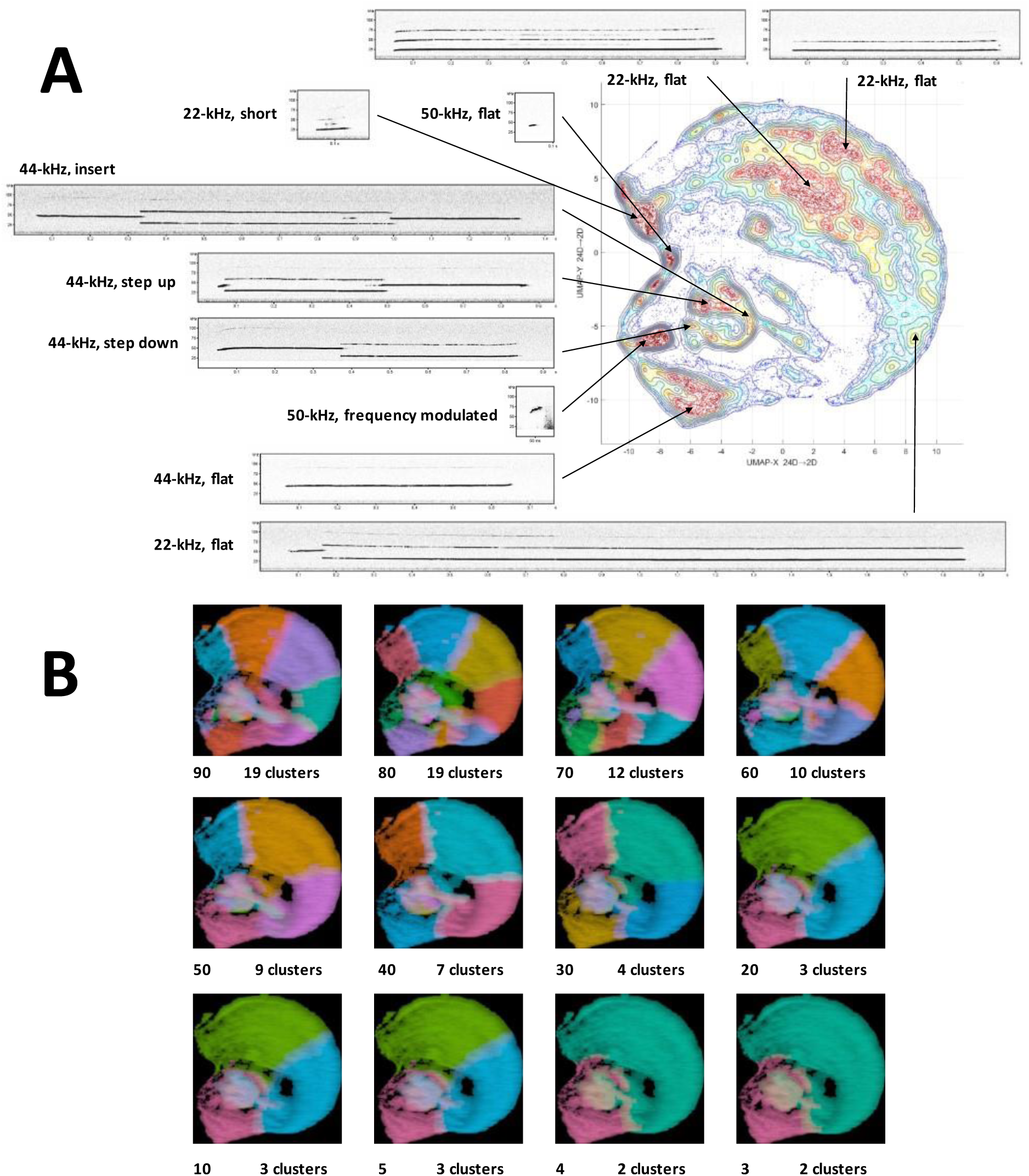
Clustering of ultrasonic vocalizations from rats emitting 44-kHz calls using UMAP projection and k-means. **A** – topological plot of ultrasonic calls using UMAP embedding from selected rats emitting 44-kHz vocalizations during trace and delay fear conditioning training (n = 26, selected from Tab. 1/Exp. 1-3/#2,4,7,8,11-13), total number of calls n = 40,084, with spectrogram miniatures pointing to the general location from which they originated. **B** – comparison of unsupervised k-means clustering with different maximum possible number of clusters using elbow optimization (different clusters denoted by colors) done by DeepSqueak software, superposed over UMAP topological plot, number on the bottom left of the miniature denotes the maximum possible number of clusters set for elbow optimization, number on the bottom right denotes the resulting number of clusters after elbow optimization.

**Fig. 3S2.**
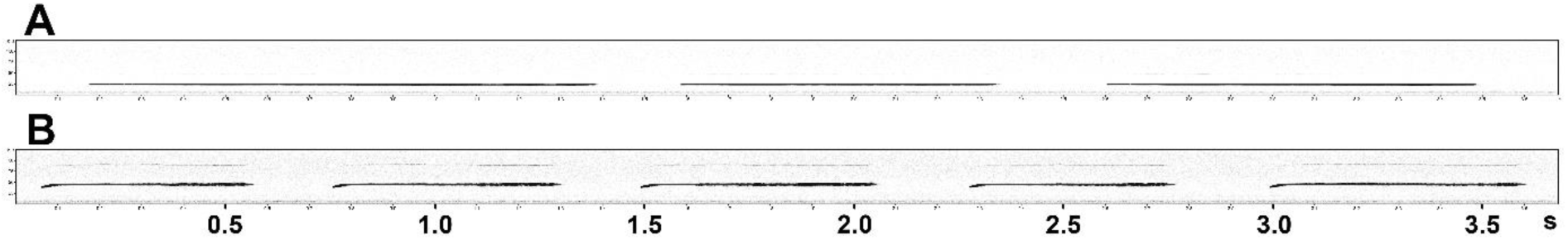
Examples of parts of ultrasonic bouts comprised of long 22-kHz vocalizations (**A**) and 44-kHz vocalizations (**B**) emitted by Wistar rats during fear-conditioning training. The average frequencies of these particular calls are 23.4 kHz (**A**) and 44.5 kHz (**B**); each vertical scale is 125 kHz.

**Fig. 4S1.**
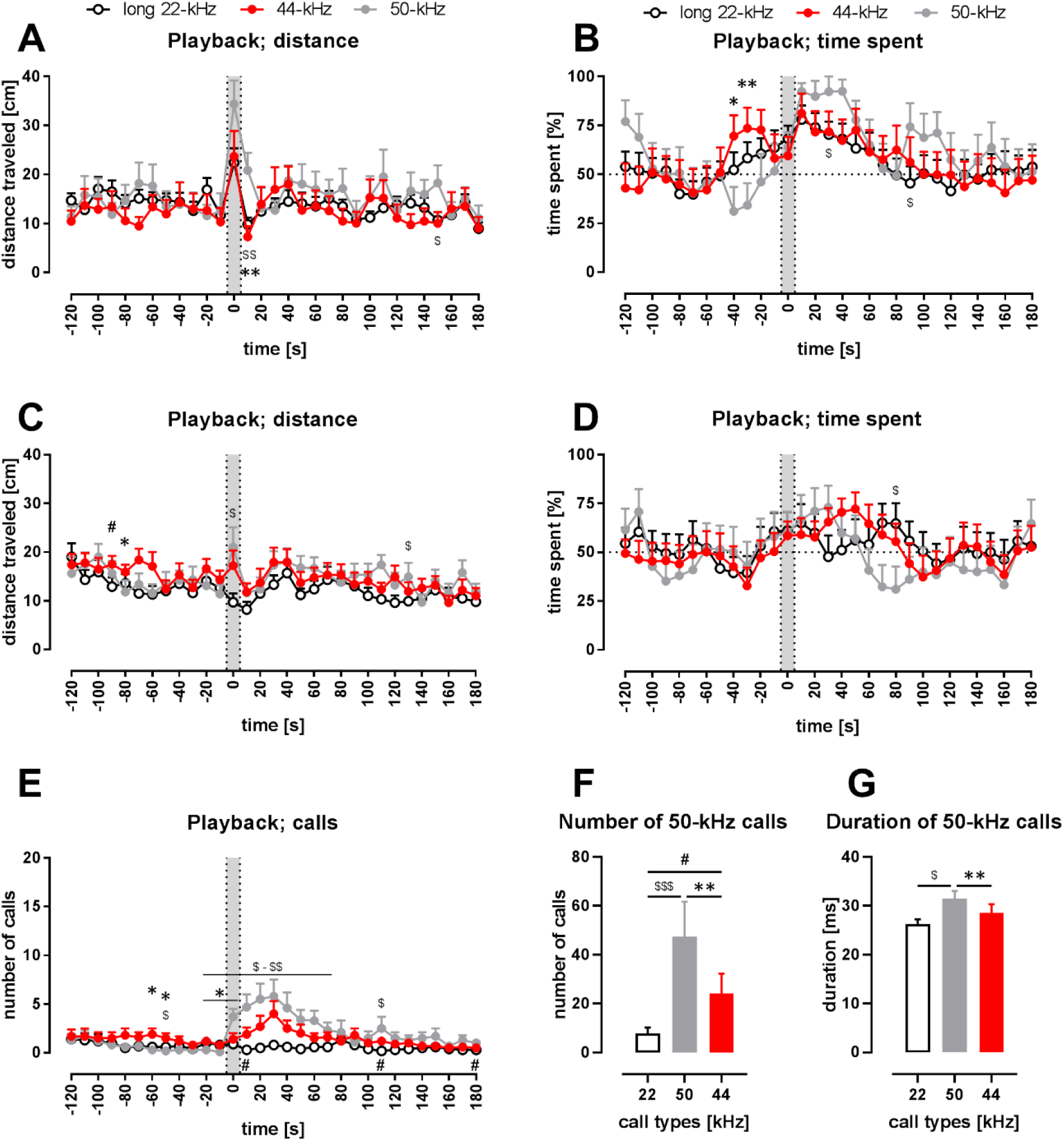
Behavioral response to playback of 44-kHz calls (vs. 50-kHz and 22-kHz calls). **AB** – rats with implanted heart-rate transmitters (comp. Fig. 4), Wistar, n = 13-16; **C-G** – rats without transmitters, Sprague-Dawley, n = 15; **AC** – distance traveled; **BD** – time spent in the speaker’s half of the cage; the dotted horizontal line marks a 50% chance value for time in a side of the cage; **E** – number of emitted vocalizations; **A-E** – gray sections correspond to the 10-s-long ultrasonic presentation, each point is a mean for a 10-s-long time-interval with SEM. **FG** – properties of 50-kHz vocalizations emitted in response to ultrasonic playback, i.e., number of calls (**F**) and duration (**G**) in 0-120 s range. **A-D** – playback presentation resulted in increased motor activity in case of, especially, 50-kHz playback and 44-kHz playback. Also, all kinds of playback resulted in increased time spent in the half of the cage next to the speaker. **E** – 50-kHz playback resulted in a rise of the number of evoked vocalizations (average from -30 to -10 time-intervals aka *before* vs. 10-30 s time-interval, p = 0.0010) as was the case with 44-kHz playback (p = 0.0142), respectively, while no rise was observed following 22-kHz playback (p = 0.2271, all Wilcoxon). However, since the increase in vocalization was robust in case of 50-kHz playback, the number of emitted vocalizations was higher than both after 22-kHz playback (e.g., p < 0.01 during 0-20 time-intervals) and after 44-kHz playback (p = 0.0172, 0 s time-interval, all Mann-Whitney). Finally, when the increases in the number of emitted ultrasonic calls in comparison with *before* intervals were analyzed, there was a difference following 44-kHz vs. 22-kHz playbacks during the 40 s time interval (p = 0.0017, Wilcoxon, comp. Fig. 4B). **F** – During the 2 min following the onset of the playbacks, the rats emitted more ultrasonic calls during and after 50-kHz playback in comparison with 22-kHz (p = 0.0002) and 44-kHz (p = 0.0067) playbacks; also, the rats emitted more ultrasonic calls during and after 44-kHz playback in comparison with 22-kHz playback (p = 0.0369), comp. Fig. 4C; all Wilcoxon). **G** – Ultrasonic 50-kHz calls emitted in response differed also in their duration, i.e., they were shorter to 22-kHz (p = 0.0195) and 44-kHz (p = 0.0039) playbacks than to 50-kHz playback. The difference between the effects of 22-kHz and 44-kHz playbacks was not significant (p = 0.5469, comp. Fig. 4D; all Wilcoxon). * 50-kHz vs. 44-kHz, $ 50-kHz vs. 22-kHz, # 22-kHz vs. 44-kHz; one character (*, $ or #), p < 0.05; two, p < 0.01; three, p < 0.001; Mann-Whitney (**AB**) or Wilcoxon (**CD**). Values are means ± SEM.

**Tab. 1.** All fear conditioning (FC) experiments described in the text. * – control groups.

| # | Exp. | FC type | strain | # of trials (shocks) | # of rats (n) | housing | transmitters | age in weeks | description/history |
| --- | --- | --- | --- | --- | --- | --- | --- | --- | --- |
| 1 | 1 | trace | Wistar | 0* | 7 | single | + | 13 | Rats that undergone FC after playback experiments (published in Olszyński et al., 2020) |
| 2 |  |  |  | 10 | 7 |  |  |  |  |
| 3 |  |  |  | 0* | 10 | paired |  |  |  |
| 4 |  |  |  | 10 | 10 |  |  |  |  |
| 5 | 2 | delay | Wistar | 0* | 37 | paired | + | 12 | FC experiments which were described before in detail (Olszyński et al., 2021, Olszyński et al., 2022) |
| 6 |  |  |  | 1 | 16 |  |  |  |  |
| 7 |  |  |  | 6 | 22 |  |  |  |  |
| 8 |  |  |  | 10 | 19 |  |  |  |  |
| 9 |  |  | SHR | 0* | 31 |  |  |  |  |
| 10 |  |  |  | 1 | 17 |  |  |  |  |
| 11 |  |  |  | 6 | 17 |  |  |  |  |
| 12 |  |  |  | 10 | 15 |  |  |  |  |
| 13 | 3 | delay | Wistar | 10 | 10 | paired | - | 12 | New FC experiment |

**Tab. 2.** Freezing associated with emission of long, monotonous vocalizations. All Wistar rats which undergone 10 trials of fear conditioning were analyzed (Tab. 1/Exp. 1-3/#2,4,8,13; n = 46). **A.** Freezing (%) in 10-s-long bins where rats emitted exclusively long 22-kHz vocalizations vs. exclusively 44-kHz vocalizations. Results were compared to baseline freezing levels before conditioning (first 5 min) and during 10-s-long periods with no vocalizations (w/o calls). More information in the text. *** vs. “first 5 min”, p < 0.001; ^#^ vs. “w/o calls”, p < 0.05; both Wilcoxon; NA, not analyzed. **B**. Freezing during the emission episodes of long 22-kHz and 44-kHz calls. Pairs of 44-kHz and long 22-kHz vocalizations were randomly selected from each animal. Freezing levels (%) did not differ between 22-kHz vs. 44-kHz calls (0.2054–0.7776 p levels, Wilcoxon). Minimum freezing duration used: 30 frames (**A**), 3 frames (for pairs of ≥ 150 ms vocalizations) or 5, 10, and 15 frames for ≥ 500 ms vocalizations (**B**).

| **A. 10-s-long time intervals; analyzed with 30 frames** |  |  |  |  |
| --- | --- | --- | --- | --- |
| group of rats analyzed | freezing [%] |  |  |  |
|  | first 5 min | 10-s bins with: |  |  |
|  |  | w/o calls | 22-kHz calls only | 44-kHz calls only |
| rats with long 22-kHz calls (n = 41) | 6.3 ± 2.3 | 39.7 ± 3.5 | 47.8 ± 3.3 ***, # | NA |
| rats with long 22-kHz calls and 44-kHz calls (n = 21) | 9.1 ± 4.2 | 33.7 ± 4.9 | 40.4 ± 4.3 *** | 50.3 ± 7.0 ***, # |
| **B. freezing behavior during selected calls; analyzed with reduced number of frames** |  |  |  |  |
| rats with long 22-kHz calls and 44-kHz calls |  | freezing [%] during emission of: |  |  |
|  | no. of frames | long 22-kHz calls | 44-kHz calls |  |
| with calls of ≥ 150 ms duration (n = 32) | 3 | 49.5 ± 7.6 | 58.8 ± 8.0 |  |
| with calls of ≥ 500 ms duration (n = 28) | 5 | 67.3 ± 7.3 | 54.1 ± 9.1 |  |
|  | 10 | 61.9 ± 8.1 | 53.3 ± 9.1 |  |
|  | 15 | 52.6 ± 9.0 | 48.4 ± 9.4 |  |

## References

Antoniadis, E. A. & Mcdonald, R. J. 1999. Discriminative fear conditioning to context expressed by multiple measures of fear in the rat. Behav Brain Res, 101, 1–13.

Avisoft Bioacoustics, CM16/CMPA ultrasound microphone specifications, Avisoft Bioacoustics website, accessed 11 October 2023.

Biały, M., Podobinska, M., Barski, J., Bogacki-Rychlik, W. & Sajdel-Sulkowska, E. M. 2019. Distinct classes of low frequency ultrasonic vocalizations in rats during sexual interactions relate to different emotional states. Acta Neurobiol Exp (Wars), 79, 1–12.

Bonauto, S. M., Greuel, O. M. & Honeycutt, J. A. 2023. Playback of rat 22-kHz ultrasonic vocalizations as a translational assay of negative affective states: An analysis of evoked behavior and brain activity. Neurosci Biobehav Rev, 153, 105396.

Bowling, D. L. & Purves, D. 2015. A biological rationale for musical consonance. Proc Natl Acad Sci U S A, 112, 11155–60.

Briefer, E. F., Padilla De La Torre, M. & Mcelligott, A. G. 2012. Mother goats do not forget their kids’ calls. Proc Biol Sci, 279, 3749–55.

Brudzynski, S. M. 2013. Ethotransmission: communication of emotional states through ultrasonic vocalization in rats. Curr Opin Neurobiol, 23, 310–7.

Brudzynski, S. M. 2019. Emission of 22 kHz vocalizations in rats as an evolutionary equivalent of human crying: Relationship to depression. Behav Brain Res, 363, 1–12.

Brudzynski, S. M. 2021. Biological Functions of Rat Ultrasonic Vocalizations, Arousal Mechanisms, and Call Initiation. Brain Sci, 11.

Brudzynski, S. M. & Bihari, F. 1990. Ultrasonic vocalization in rats produced by cholinergic stimulation of the brain. Neurosci Lett, 109, 222–6.

Brudzynski, S. M., Bihari, F., Ociepa, D. & Fu, X. W. 1993. Analysis of 22 kHz ultrasonic vocalization in laboratory rats: long and short calls. Physiol Behav, 54, 215–21.

Coffey, K. R., Marx, R. G. & Neumaier, J. F. 2019. DeepSqueak: a deep learning-based system for detection and analysis of ultrasonic vocalizations. Neuropsychopharmacology, 44, 859–868.

Darwin, C. 1872. “Chapter 4: Means of Expression in Animals”, The Expression of the Emotions in Man and Animals, New York, D. Appleton & Company.

Dos Santos Correa, M., Vaz, B. D. S., Grisanti, G. D. V., De Paiva, J. P. Q., Tiba, P. A. & Fornari, R. V. 2019. Relationship between footshock intensity, post-training corticosterone release and contextual fear memory specificity over time. Psychoneuroendocrinology, 110, 104447.

Dupin, M., Garcia, S., Boulanger-Bertolus, J., Buonviso, N. & Mouly, A. M. 2019. New Insights from 22-kHz Ultrasonic Vocalizations to Characterize Fear Responses: Relationship with Respiration and Brain Oscillatory Dynamics. eNeuro, 6.

Ester, M., Kriegel, H.-P., Sander, J. & Xu, X. A density-based algorithm for discovering clusters in large spatial databases with noise. kdd, 1996. 226–231.

Fanselow, M. S. & Bolles, R. C. 1979. Naloxone and shock-elicited freezing in the rat. J Comp Physiol Psychol, 93, 736–44.

Finnie, P. S. B. & Nader, K. 2020. Amyloid Beta Secreted during Consolidation Prevents Memory Malleability. Curr Biol, 30, 1934–1940 e4.

Garcia, E. J., Mccowan, T. J. & Cain, M. E. 2015. Harmonic and frequency modulated ultrasonic vocalizations reveal differences in conditioned and unconditioned reward processing. Behav Brain Res, 287, 207–14.

Gonzalez-Palomares, E., Boulanger-Bertolus, J., Dupin, M., Mouly, A. M. & Hechavarria, J. C. 2023. Amplitude modulation pattern of rat distress vocalisations during fear conditioning. Sci Rep, 13, 11173.

Hakansson, J., Jiang, W., Xue, Q., Zheng, X., Ding, M., Agarwal, A. A. & Elemans, C. P. H. 2022. Aerodynamics and motor control of ultrasonic vocalizations for social communication in mice and rats. Bmc Biol, 20, 3.

Haubrich, J., Bernabo, M. & Nader, K. 2020. Noradrenergic projections from the locus coeruleus to the amygdala constrain fear memory reconsolidation. Elife, 9.

Haubrich, J. & Nader, K. 2023. Network-level changes in the brain underlie fear memory strength. Elife, 12.

Hoeschele, M. 2017. Animal Pitch Perception: Melodies and Harmonies. Comp Cogn Behav Rev, 12, 5–18.

Jahołkowski, P., Kiryk, A., Jedynak, P., Ben Abdallah, N. M., Knapska, E., Kowalczyk, A., Piechal, A., Blecharz-Klin, K., Figiel, I., Lioudyno, V., Widy-Tyszkiewicz, E., Wilczyński, G. M., Lipp, H. P., Kaczmarek, L. & Filipkowski, R. K. 2009. New hippocampal neurons are not obligatory for memory formation; cyclin D2 knockout mice with no adult brain neurogenesis show learning. Learn Mem, 16, 439–51.

Kalamari, A., Kentrop, J., Hinna Danesi, C., Graat, E. A. M., Van, I. M. H., Bakermans-Kranenburg, M. J., Joels, M. & Van Der Veen, R. 2021. Complex Housing, but Not Maternal Deprivation Affects Motivation to Liberate a Trapped Cage-Mate in an Operant Rat Task. Front Behav Neurosci, 15, 698501.

Kitch, J. A. & Oates, J. 1994. The perceptual features of vocal fatigue as self-reported by a group of actors and singers. J Voice, 8, 207–14.

Mcinnes, L., Healy, J. & Melville, J. 2018. Umap: Uniform Manifold Approximation and Projection. Journal of Open Source Software, 3(29), 861, 10.21105/joss.00861

Okamoto, K. & Aoki, K. 1963. Development of a strain of spontaneously hypertensive rats. Jpn Circ J, 27, 282–93.

Olszyński, K. H., Polowy, R., Małż, M., Boguszewski, P. M. & Filipkowski, R. K. 2020. Playback of Alarm and Appetitive Calls Differentially Impacts Vocal, Heart-Rate, and Motor Response in Rats. iScience, 23, 101577.

Olszyński, K. H., Polowy, R., Wardak, A. D., Grymanowska, A. W. & Filipkowski, R. K. 2021. Increased Vocalization of Rats in Response to Ultrasonic Playback as a Sign of Hypervigilance Following Fear Conditioning. Brain Sci, 11.

Olszyński, K. H., Polowy, R., Wardak, A. D., Grymanowska, A. W., Zieliński, J. & Filipkowski, R. K. 2022. Spontaneously hypertensive rats manifest deficits in emotional response to 22-kHz and 50-kHz ultrasonic playback. Prog Neuropsychopharmacol Biol Psychiatry, 120, 110615.

Pestana-Oliveira, N., Nahey, D. B., Johnson, T. & Collister, J. P. 2020. Development of the Deoxycorticosterone Acetate (Doca)-salt Hypertensive Rat Model. Bio Protoc, 10, e3708.

Potasiewicz, A., Holuj, M., Litwa, E., Gzielo, K., Socha, L., Popik, P. & Nikiforuk, A. 2020. Social dysfunction in the neurodevelopmental model of schizophrenia in male and female rats: Behavioural and biochemical studies. Neuropharmacology, 170, 108040.

Poulos, A. M., Mehta, N., Lu, B., Amir, D., Livingston, B., Santarelli, A., Zhuravka, I. & Fanselow, M. S. 2016. Conditioning- and time-dependent increases in context fear and generalization. Learn Mem, 23, 379–85.

Riede, T. 2013. Stereotypic laryngeal and respiratory motor patterns generate different call types in rat ultrasound vocalization. J Exp Zool A Ecol Genet Physiol, 319, 213–24.

Rousseeuw, P. J. 1987. Silhouettes: a graphical aid to the interpretation and validation of cluster analysis. Journal of computational and applied mathematics, 20, 53–65.

Saito, Y., Tachibana, R. O. & Okanoya, K. 2019. Acoustical cues for perception of emotional vocalizations in rats. Sci Rep, 9, 10539.

Sales, G. D. 1979. Strain Differences in the Ultrasonic Behavior of Rats (Rattus-Norvegicus). American Zoologist, 19, 513–527.

Sales, G. D. 1991. The effect of 22 kHz calls and artificial 38 kHz signals on activity in rats. Behav Processes, 24, 83–93.

Shimoju, R., Shibata, H., Hori, M. & Kurosawa, M. 2020. Stroking stimulation of the skin elicits 50-kHz ultrasonic vocalizations in young adult rats. Journal of Physiological Sciences, 70.

Simola, N. & Granon, S. 2019. Ultrasonic vocalizations as a tool in studying emotional states in rodent models of social behavior and brain disease. Neuropharmacology, 159, 107420.

Silkstone, M. & Brudzynski, S. M. 2019. The antagonistic relationship between aversive and appetitive emotional states in rats as studied by pharmacologically-induced ultrasonic vocalization from the nucleus accumbens and lateral septum. Pharmacol Biochem Behav, 181, 77–85.

Sonninen, A. & Hurme, P. 1998. Vocal fold strain and vocal pitch in singing: radiographic observations of singers and nonsingers. J Voice, 12, 274–86.

Taylor, J. O., Urbano, C. M. & Cooper, B. G. 2017. Differential patterns of constant frequency 50 and 22 khz usv production are related to intensity of negative affective state. Behav Neurosci, 131, 115–126.

Turner, C. A., Hagenauer, M. H., Aurbach, E. L., Maras, P. M., Fournier, C. L., Blandino, P., Jr., Chauhan, R. B., Panksepp, J., Watson, S. J., Jr. & Akil, H. 2019. Effects of early-life FGF2 on ultrasonic vocalizations (USVs) and the mu-opioid receptor in male Sprague-Dawley rats selectively-bred for differences in their response to novelty. Brain Res, 1715, 106–114.

Wang, S. H., De Oliveira Alvares, L. & Nader, K. 2009. Cellular and systems mechanisms of memory strength as a constraint on auditory fear reconsolidation. Nat Neurosci, 12, 905–12.

Wöhr, M., Borta, A. & Schwarting, R. K. 2005. Overt behavior and ultrasonic vocalization in a fear conditioning paradigm: a dose-response study in the rat. Neurobiol Learn Mem, 84, 228–40.

